# Loss-of-function mutation in the polyamine transporter gene *OsLAT5* as a selectable marker for genome editing

**DOI:** 10.1101/2023.12.12.571390

**Authors:** Kyrylo Schenstnyi, Zhengzhi Zhang, Bo Liu, Masayoshi Nakamura, Van Schepler-Luu, Eliza P.I. Loo, Bing Yang, Wolf B. Frommer

## Abstract

Genome editing by TALENs, CRISPR/Cas, base or prime editing have become routine tools. During stable plant transformation, the gene coding for the editing enzyme, e.g., *Cas9,* the guide RNAs (gRNAs), alongside a selectable marker are integrated into the nuclear genome. Identification of successful transformants relies on selectable or screenable markers, typically genes providing resistance to antibiotics or herbicides. Selectable markers use a substantial portion of the T-DNA, hence reducing transfer efficiency by limiting the effective number of TALENs or guide/pegRNAs that can be used in parallel. Moreover, marker genes are frequently subject to gene silencing. Here, we generated loss-of-function mutations in PUT/LAT-type polyamine transporter family genes to confer resistance to the phytotoxin methylviologen (MV) as a method for selection. As a proof of concept, CRISPR/Cas9 vectors with gRNAs were constructed to target three close homologs *OsLAT1*, *OsLAT5*, and *OsLAT7*. We show that loss of *OsLAT5* (also known as *OsPUT3* or *OsPAR1*) function was sufficient to confer resistance to MV in rice seeds, seedlings and calli, validating the editing approach of *OsLAT5* to obtain a selectable marker. We discuss the potential of incorporating a gRNA cassette (for *OsLAT5*) as a selectable marker and a reporter for successful genome editing for optimizing editing protocols.

## Introduction

Genome editing is widely used in both model plant and crop species for research and biotechnology [1,2]. Combinations of CRISPR/Cas9/Cpf1 and prime editing has successfully generated elite rice varieties with broad-spectrum resistance to bacterial blight and other diseases [3–9]. Major advantages of genome editing over classical breeding include the precision of prime editing, speed, and the ability to target multiple traits simultaneously without linkage drag. At present, most genome editing approaches in plants require stable transformation with a CRISPR construct. Due to the low transformation efficiencies, screenable or selectable markers have to be implemented on the same construct [10,11]. A selectable marker typically comprises a cassette consisting of a strong ubiquitous plant promoter, an antibiotic or herbicide resistance gene, and a terminator. Selectable marker cassettes are comparatively large, using up a substantial fraction of the capacity of the *Agrobacterium* T-DNA. The capacity/insert size of a typical bacterial plasmid is 5-25 kb; insertion into the host chromosome is initiated from the right border, but there is a length-dependent decrease in efficiency of the insertion of sequences towards the left border [12](ref). When the size of the components necessary for replication and selection in bacteria, and the size of Cas9/Cpf1 modules are taken into account, the number of gRNAs that can be included into a single CRISPR construct, i.e., for multiplex genome editing, is limited by the size of a selectable marker for *in planta* selection. To fully utilize the available nucleotides for gRNAs, we hypothesize that a selectable marker can be created by editing the host plant’s endogenous trait. Ideally, loss of a certain gene function that would lead to acquisition of a new trait, i.e., resistance to a chemical, metabolite or xenobiotic. The gain of a new trait also serves as a control for successful editing. Such loss-of-function markers generated by editing would eliminate the need to remove the herbicide marker, and could be used also for transgene-free genome editing approaches [13–15]. To avoid potential issues associated with *Agrobacterium*-mediated T-DNA delivery, chemically synthesized gRNA and purified Cas proteins pre-assembled into ribonucleoproteins (RNPs) can be used for transient transgene-free genome editing in plant cells [16,17]. Despite the availability of studies reporting successful transgene-free genome editing, editing efficiencies remain too low to edit without selection [18–21].

A member of the *Arabidopsis* L-type amino acid transporter (LAT) family transporter (SLC7A5), named RMV1 (<u>R</u>esistant to <u>M</u>ethyl <u>V</u>iologen 1) was identified to be involved in the transmembrane transport of polyamines (PA) and N,N′-dimethyl-4,4′-bipyridinium dichloride methylviologen (MV; trademark paraquat; PQ)[22,23]. Genes encoding L-type amino acid transporters were identified multiple times independently and named *RMV*, *LAT*, *PUT* (*<u>P</u>olyamine <u>U</u>ptake <u>T</u>ransporter*), *LHR* (*<u>L</u>ower expression of <u>H</u>eat-<u>r</u>esponsive <u>G</u>ene*) or *PAR* (*<u>Pa</u>raquat <u>r</u>esistant*) [24]. The rice genome contains nine *LAT*/*PUT*/*PAR* paralogs [22,23]. When expressed in protoplasts of *Arabidopsis* or rice, OsPAR1-GFP fusion protein colocalized with the Golgi marker GmMan1-mCherry [25]. Following the initial uptake of MV by plasma membrane-localized PUT/LAT-type transporters, i.e., *OsLAT1/OsPUT1* and rice ortholog of AtPDR11/AtPQT24 [22,23,26,27], the Golgi-localized OsLAT5/OsPUT3/OsPAR1 transfers MV to the cytosol, hence making it available to chloroplasts, [23,25,28]. RNAi of *OsLAT5/OsPUT3/OsPAR1* caused partial MV resistance resulting into reduction of chlorophyll content (≈ 20-35%) when plants were treated with the relatively high concentrations of MV (5 μM). RNAi lines had a normal grain weight and did not display detectable phenotypic differences compared to wild type controls even in field growth conditions indicating that knockdown of *OsLAT5/OsPUT3/OsPAR1* has no detrimental effects on the growth and development of rice [25].

Based on the partial MV resistance of *OsLAT5/OsPUT3/OsPAR1* RNAi rice lines, we hypothesized that PUT/LAT-type transporter-encoding genes could be suitable selectable markers for genome editing approaches.. We generated individual mutants in three paralogous genes, i.e., *OsLAT1/OsPUT1*, *OsLAT5/OsPUT3/OsPAR1* and *OsLAT7/OsPUT2*, and found that a loss-of-function mutation in *OsLAT5/OsPUT3/OsPAR1* is sufficient to enable selection in axenic cultures during callus induction and regeneration of plants from calli as well as selection of T1 plants during germination. Notably, loss-of-function mutations in the two close paralogs *OsLAT1/OsPUT1* and *OsLAT7/OsPUT2* did not convey resistance under these conditions. An independent study performed in parallel confirmed that loss-of-function mutations in *AtPAR1* generated by CRIPSR/Cas9 can be used to identify edited lines [14]. A similar approach that targeted the same three paralogs in rice as targeted here using a single construct showed that the triple *oslat5/osput3/ospar1, oslat1/osput1, oslat7/osput2* mutants were also resistant to MV [29]. Neither the single mutants characterized here, nor the triple mutants showed apparent growth or developmental defects under normal growth conditions [29]. We therefore surmise that the generation of loss-of-function mutations by editing in the single *OsLAT5/OsPUT3/OsPAR1* locus is sufficient to obtain MV resistance, and we show that MV can be used as a selectable marker during callus induction and regeneration of plants from calli to identify transformants or in the seedling stage to detect lines carrying the edited locus.

## Materials and methods

### Sequence alignments and phylogenetic analysis

Accession numbers are: *OsLAT1*/*OsPUT1* (Os02g0700500; LOC_Os02g47210), *OsLAT5*/*OsPUT3*/*OsPAR1* (Os03g0576900; LOC_Os03g37984), *OsLAT7*/*OsPUT2* (Os12g0580400; LOC_Os12g39080). and *AtPAR1* (At1G31830; NCBI accession number NM_102920). Phylogenetic trees were built using Geneious Tree Builder in Geneious Prime 2023.1.1 software with standard settings. Protein sequences were extracted from UniProt (AtPAR1, Q9C6S5; OsLAT1/OsPUT1, Q6Z8D0; OsLAT5/OsPUT3/OsPAR1, Q10HT5; OsLAT7/OsPUT2, A0A0N7KU97). Sequences encoding transmembrane domains were highlighted according to the annotation from UniProt. Alignments of nucleotide sequences were made using MUSCLE (<u>Mu</u>ltiple <u>S</u>equence <u>C</u>omparison by <u>L</u>og-<u>E</u>xpectation), while alignments of amino acid sequences were generated with Clustal Omega. Nucleotides or amino acids that are conserved in the majority of the analyzed sequences are shaded in BLACK using pyBoxshade program.

### Rice cultivation

Rice seed germination and plant cultivation were done as described [30]. Briefly, rice seeds were sterilized and germinated on ½-salt strength MS medium (2.2 grams / liter of Murashige & Skoog basal salt mixture including vitamins from Duchefa Biochemie; Catalog # M0222) supplemented with 1% sucrose (Duchefa Biochemie; Product # S0809). Seedlings were grown for 10 days in tissue culture vessels (GA-7 Vessel, Magenta LLC) before transfer to soil. Plants were grown in greenhouses maintained at 30 °C day / 25 °C night, relative humidity (RH) 50-70% with supplemental LED light (Valoya, BX100 NS1) using a 8h light / 16h dark photoperiod (400 μmol/m^-2^s^-1^). Plants were fertilized weekly from the 2^nd^ week and biweekly from the 6^th^ week after germination with Peters Excel CalMag Grower 15:5:15+7CaO+3MgO+TE fertilizer (ICL Specialty Fertilizers).

### CRISPR constructs

Four vectors were used to generate three CRISPR-Cas9 constructs targeting *OsLAT1*, *OsLAT5*, and *OsLAT7*, respectively. Modular intermediate vectors ptGgRNA1 and ptGRNA-T2 were used to make two guides. The recipient vector pENTR4-U6.1P-ccdBchl was used to assemble the gRNA units. Subsequently the assembly was mobilized into the binary Gateway vector pBY02rCas9-GW [31]. To target *OsLAT5*, one pair of complementary oligos with different four-base overhangs at the 5ߣ-ends (gLAT5-F1 and gLAT5-R1) were denatured and annealed to form a double-stranded DNA fragment (dsOligo), which was cloned into ptGRNA1 at *BsmB*I sites. Similarly, a pair of oligonucleotides (gLAT5-F2 and gLAT5-R2) were cloned into ptGRNA-T2. After confirmation of successful insertion of the dsOligos, the two resulting plasmids were used along with pENTR4-U6.1P-ccdBchl to perform Golden Gate reactions using *Bsa*I and T4 ligase, resulting in substitution of *ccdBchl* cassette by the two gRNA units (tRNA-gRNA architecture) under control of rice *U6* promoter (U6.1P). The tRNA-gRNA cassette, flanked by *att*L1-*att*L2, was integrated into pBY02rCas9-GW via LR reactions (Invitrogen Gateway™ LR Clonase™ II Enzyme mix; Catalog # 11791020). The final plasmid (pBY02rCas9_gLAT5) was restricted with *BamH*I, *Hind*III to validate the correct insert size. Using the same approach but with different oligo pairs, CRISPR constructs pBY02rCas9_gLAT1 and pBY02rCas9_gLAT7 targeting *OsLAT1* and *OsLAT7*, respectively, were generated. Sequences of oligonucleotides are provided in S1. <u>Stable rice transformation</u> The rice cultivar Kitaake (*Oryza sativa* subsp. *japonica*) was used for *Agrobacterium*-mediated transformation using the method published by Wang et al., 2017 [32]. Briefly, *Agrobacterium* strain EHA105 carrying a CRISPR construct was used to infect embryo-derived rice calli for 3 days. The calli were selected on callus-inducing medium supplemented with Hygromycin B (50 mg/L; Sigma Aldrich) for two rounds (14 days each round). Hygromycin-resistant calli were cultured on regeneration medium for 1 or 2 rounds (2 weeks per round) for shoot initiation. Shoots were moved to rooting medium for root initiation and elongation before transferring to soil and were grown in the greenhouse at 28 °C /12h light and 24°C / 12h dark photoperiod.

### Genotyping of edited rice lines

Since the gRNA target sites / Cas9 cleavage sites were designed to overlap with cleavage sites of restriction enzymes therefore, introduction of mutations by Cas9 results in loss of restriction sites. To detect mutations, T0 and successive generation plants were sampled for DNA extraction using CTAB method [33]. Gene-specific primers were used for amplification of relevant genomic fragments via polymerase chain reaction (PCR). PCR-derived amplicons were restricted to identify the amplicons that lost restriction sites. Sanger sequencing of PCR products was used to determine mutations. Plantlets carrying mutations in target genes were grown to maturity for seed multiplication and further selected for homozygosity of targeted loci. Sequences of oligonucleotides used for genotyping are provided in S1.

### MV selection methods for seedlings and calli

Seeds were surface-sterilized with 70% ethanol for one minute, washed in 6.5 % sodium hypochlorite (NaOCl) solution for 10 minutes, and washed three times with autoclaved MilliQ water. For evaluation of MV sensitivity during seed germination, seeds were sown on ½-salt strength MS medium (2.2 grams / liter of Murashige & Skoog basal salt mixture including vitamins from Duchefa Biochemie; Catalog # M0222) supplemented with 1% sucrose (Duchefa Biochemie; Catalog # S0809), and either containing dimethyl sulfoxide (DMSO; C2H6OS, Fisher Chemical; Catalog # D/4121/PB15) as a solvent, or methyl viologen dichloride hydrate (C12H14Cl2N2 x H2O, Sigma-Aldrich; Catalog # 856177-1G). Nine days after sowing on ½-salt strength MS medium, seedling shoot length was measured with a metric ruler. For evaluation of MV sensitivity during seedling transfer, seeds were sown on ½-salt strength MS medium supplemented with 1% sucrose. Four days after seed sowing, seedlings were transferred to ½-salt strength MS medium supplemented with 1% sucrose, either containing solvent (DMSO), or methyl viologen dichloride hydrate. Five days after the transfer, seedling shoot length was measured with a metric ruler. For evaluation of MV sensitivity during callus induction, seeds were sown on callus-inducing medium. After 10 days calli started to emerge. Calli were transferred to the fresh callus-inducing medium three times in the intervals of two weeks. Callus-inducing medium recipe was as described [32].

### Statistical analyses

If not stated otherwise, graphical representations and statistical analyses were prepared using R 4.3.0 on RStudio 2021.09.2. Data were drawn as boxplots using package ggplot2 (https://ggplot2.tidyverse.org). The boxplot is delimited by the first and the third quartile of the distribution of the studied variable. The line inside the boxplot represents the median of the variable. Finally, the two lines that start from the boxplot join the minimum and maximum theoretical values. Each recorded datapoint is represented by the black or red dot. Red dots show the recorded datapoint for a seedling that was used for a photo. The total number (*n*) of biological samples for each genotype and condition equals 16. Statistics were calculated using Pairwise Wilcoxon Rank Sum Test with the p-value correction for multiple testing using Bonferroni adjustment method.

## Results

### Three close *AtLAT1* homologs in the rice genome

The rice genome contains nine PUT/LAT-type transporter-encoding genes [30]. Phylogenetic analyses indicate that rice *LAT1/PUT1, LAT5/PUT3/PAR1*, and *LAT7/PUT2* form a separate subclade. Among these three paralogs, *OsLAT5/OsPUT3/OsPAR1* is the closest homologue of *AtPAR1* in rice (Fig 1, S2 and S3). *OsLAT5/OsPUT3/OsPAR1* is expressed in multiple rice tissues [35](S4).

**Fig 1.**
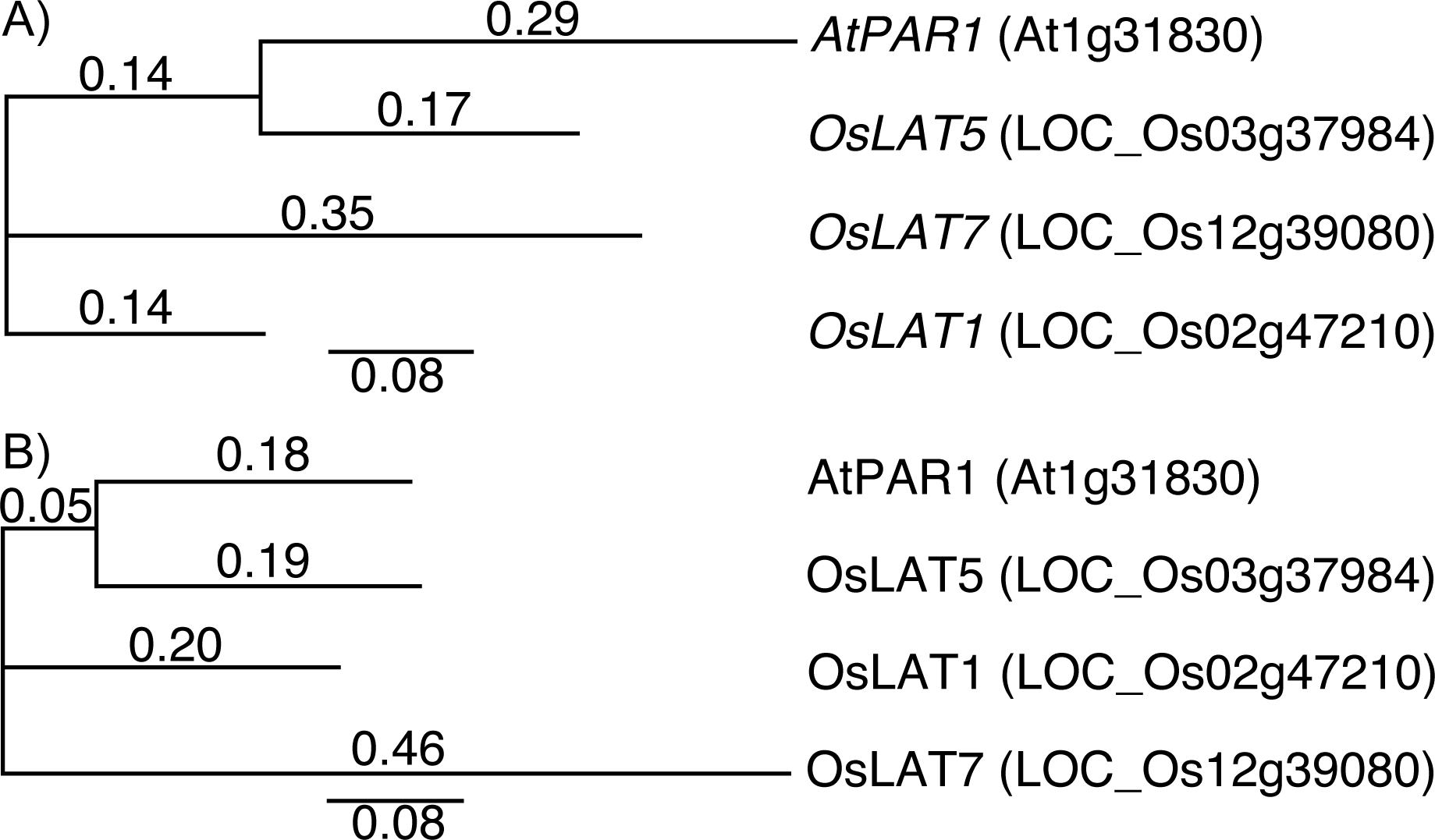
Rice *LAT1*, *LAT5,* and *LAT7* are highly homologous to *AtPAR1*. A) Neighbor-joining phylogenetic analysis of protein coding sequences of *AtPAR1* and *OsLAT1*, *OsLAT5*, and *OsLAT7*. Values on tree branches represent substitutions per site. B) Neighbor-joining phylogenetic analysis of AtPAR1 and OsLAT1, OsLAT5, and OsLAT7 proteins. Values on tree branches represent substitutions per site.

### Independent editing of three *LAT1* paralogs in rice

To evaluate whether loss-of-function mutations in one of three closely related *LAT* genes could be used for selection during rice transformation, *O. sativa* cv. Kitaake was transformed using *Agrobacterium*-mediated transformation with T-DNA constructs containing two gRNAs targeting either *LAT1/PUT1, LAT5/PUT3/PAR1* or *LAT7/PUT2*, respectively. The gRNAs target sequences were designed to lead to disruption of transmembrane spanning domains (Fig 2 and S5). Target sites in the transmembrane domains were chosen instead of the start codon, since disruption of transmembrane domains typically leads to loss of function, and editing of the start codon, in certain cases, will not cause loss of function due to the presence of alternative downstream translation start sites [36]. Two independent mutant alleles per each target gene were identified and used for further experimentation (Fig 2 and Table 1). Mutations at target genes and putative protein sequences of the edited sequences are listed in S5 and S6.

**Fig 2.**
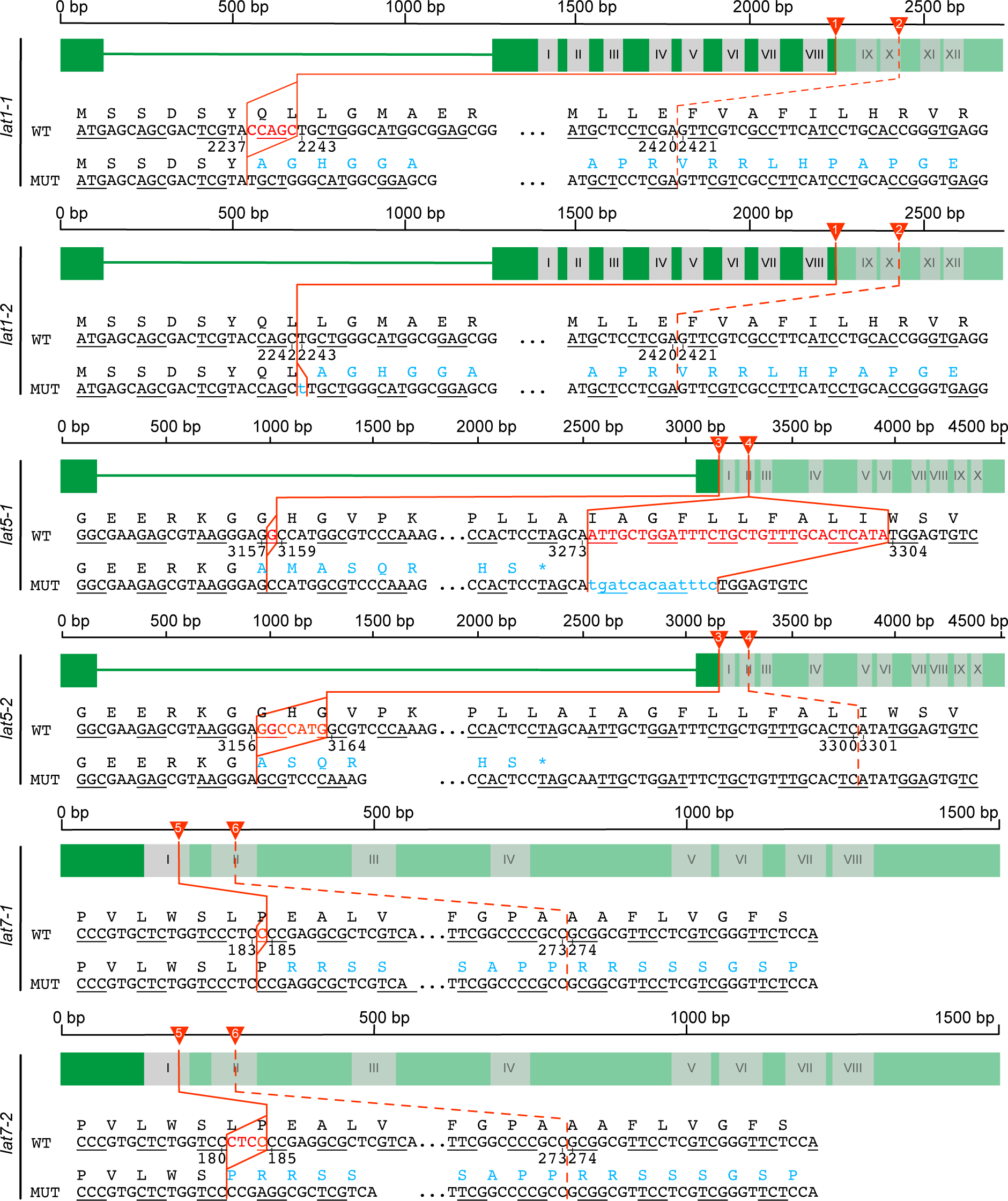
CRISPR/Cas9-mediated mutagenesis yields null *LAT1*, *LAT5*, and *LAT5* alleles. Graphical display of the CRISPR/Cas9-induced null *LAT1*, *LAT5*, and *LAT5* alleles. Black lines over green blocks provide information about the length of genomic sequences of each WT allele in base pairs (bp). Wide green horizontal bars indicate exons while green horizontal lines indicate introns. Grey shaded blocks with Roman numbers represent transmembrane domains according to the annotations from UniProt database. Faded colors indicate regions of a given mutant allele that are translated differently than a corresponding WT allele. Numbered red triangles indicate sequence parts targeted by gRNAs that were used to introduce mutations. Straight red lines highlight sequence parts that were mutated. While dash red lines represent sequence parts that were targeted by a gRNA, but were not mutated. The red font indicates deletions (upper case letters) and while insertions are highlighted with the blue font (lower case letters). Blue color indicates amino acids or translational stop codons (asterisk) that are changed due to frameshifts. Details on complete nucleotide and amino acid sequences of mutant alleles are provided in S5 and S6.

**Table 1.** Summary of loss-of-function mutations in *LAT1*, *LAT5*, and *LAT7* genes used in this study.

| Allele | Type of mutations | Generation used for MV screening |
| --- | --- | --- |
| *lat1-1* | Single site mutation: 5 bp deletion; frameshift | T2 |
| *lat1-2* | Single site mutation: 1 bp insertion; frameshift | T3 |
| *lat5-1* | Mutations at two sites: 1 bp deletion and 30 bp indel; frameshift | T4 |
| *lat5-2* | Single site mutation: 7 bp deletion; frameshift | T2 |
| *lat7-1* | Single site mutation: 1 bp deletion; frameshift | T3 |
| *lat7-2* | Single site mutation: 4 bp deletion; frameshift | T3 |

### MV resistance of seedlings carrying *LAT5* null alleles

To evaluate the effect of the frameshift mutations in the three *LAT* genes on MV resistance, seeds of homozygous mutant lines were germinated on ½-salt strength MS medium supplemented either with the solvent (0 μM MV) or 0.05 μM MV. WT, *lat1*, and *lat7* seeds were equally sensitive to MV and no statistically significant differences in sensitivity to MV between these genotypes were detected (Fig 3). Only *lat5-1* and *lat5-2* seeds germinated on MV-containing medium indicating that a loss-of-function mutation in *LAT5* is sufficient to provide resistance to MV.

**Fig 3.**
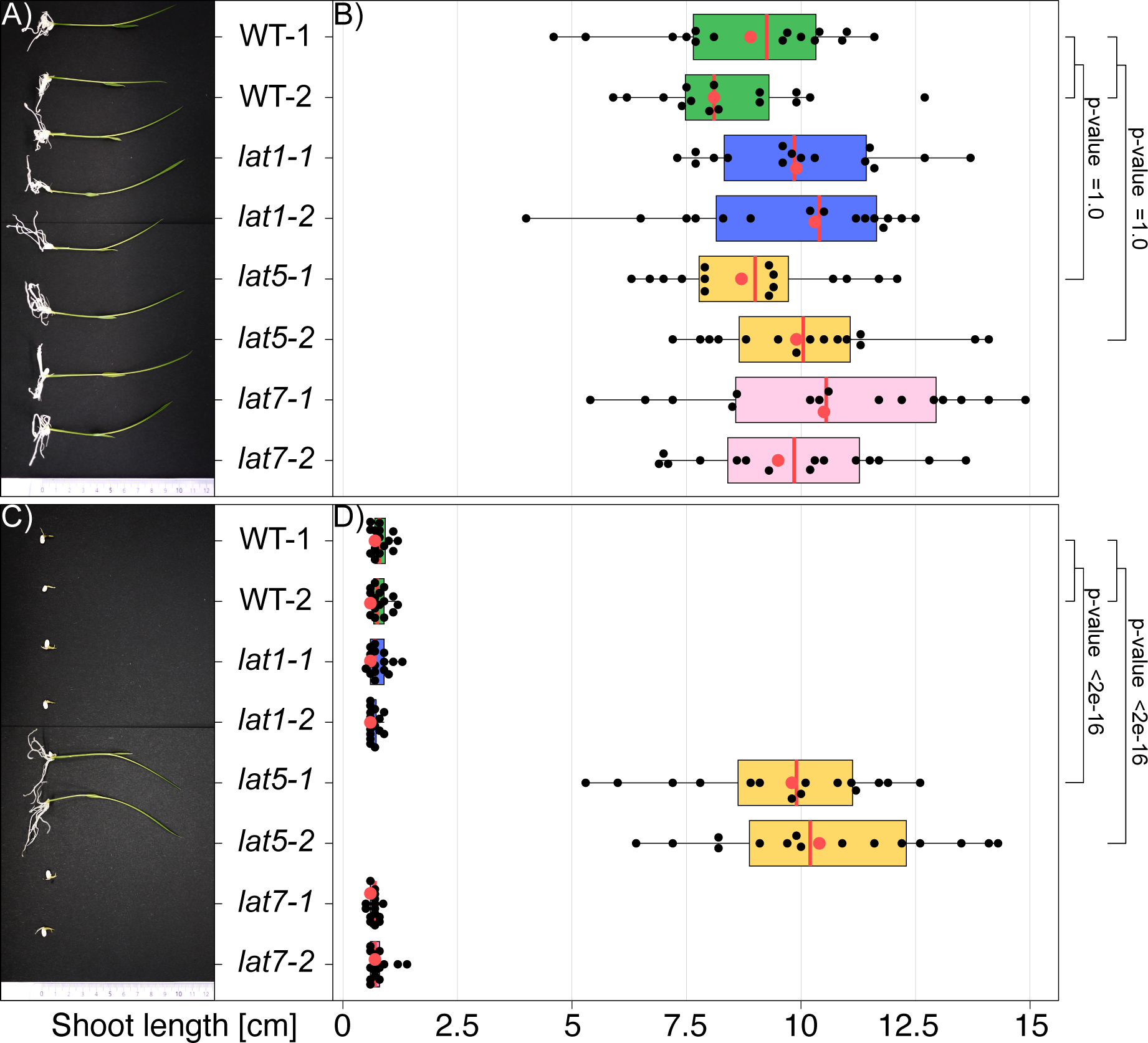
Loss-of-function mutations in *LAT5* lead to MV resistance. A) WT and loss-of-function *lat* seedlings germinated on medium lacking MV (0 μM). B) Quantification of MV resistance of WT and loss-of-function *lat* seedlings germinated on medium lacking MV (0 μM). C) WT and loss-of-function *lat* seedlings germinated on medium containing MV (0.05 μM). D) Quantification of MV resistance of WT and loss-of-function *lat* seedlings germinated on medium containing MV (0.05 μM). WT-1 and WT-2 are two independent lines of *O. sativa* cv. Kitaake harvested in 2022 and 2023, respectively. Each mutant genotype, i.e., *lat1*, *lat5*, and *lat7*, is represented by two independent lines homozygous for respective mutations (Fig 2 and Table 1). *n* = 16, where *n* is a number of seedlings representing each genotype germinated on each medium. All dots represent values for “Shoot length, cm” for individual plantlets, while red dots represent these values for plantlets that were chosen for pictures in A) and C). Box plots: red vertical line is median; box limits are lower and upper quartiles; whiskers are highest and lowest data points. Pairwise Wilcoxon Rank Sum Test with the p-value correction for multiple testing using Bonferroni adjustment method was used to calculate significant differences between groups.

Since a loss-of-function mutation in *LAT5* was sufficient to provide MV resistance, MV-based selection of primary transformants carrying null *LAT5* alleles after transgenic and transgene-free genome editing might be applicable also during shoot regeneration process. While the calli at this stage are not green, MV had been shown to also be effective in dark conditions [37]. To test this hypothesis, WT and *lat5* seeds were sowed on ½-salt strength MS medium. After four days, WT and *lat5* seedlings were transferred onto ½-salt strength MS medium supplemented either with the solvent (0 μM MV) or 0.1 μM MV. Five days after transfer only lines that contained null *lat5* alleles, namely *lat5-1* and *lat5-2*, continued development on MV-containing medium, while growth of WT seedlings was inhibited (Fig 4). These data indicate that loss-of-function mutations in *LAT5* can be used for selection of primary transformants/edited lines during shoot regeneration stage either after transgenic or transgene-free genome editing.

**Fig 4.**
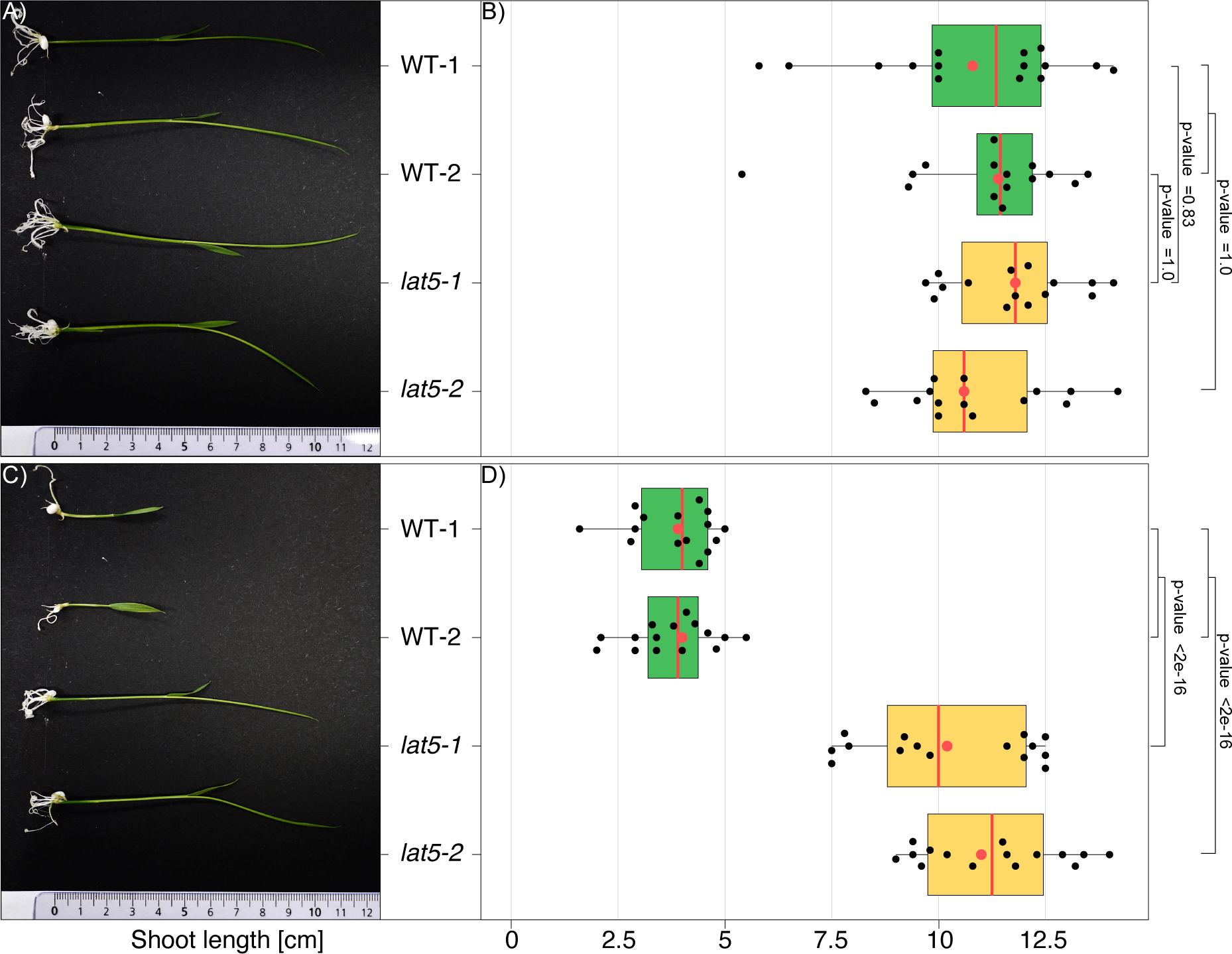
Shoots carrying null *LAT5* alleles regenerate on MV-containing medium. A) WT and loss-of-function *lat5* seedlings regenerated on medium lacking MV (0 μM). B) Quantification of seedling length of WT and loss-of-function *lat5* seedlings regenerated on medium lacking MV (0 μM). C) WT and loss-of-function *lat5* seedlings regenerated on medium containing MV (0.1 μM). D) Quantification of seedling length of WT and loss-of-function *lat5* seedlings regenerated on medium containing MV (0.1 μM). WT-1 and WT-2 are two independent lines of *O. sativa* cv. Kitaake harvested in 2022 and 2023, respectively. *lat5-1* and *lat5-2* are two independent lines homozygous for respective loss-of-function mutations in *LAT5* (Fig 2 and Table 1). Shoot length (cm) for 16 seedlings per genotype / treatment was quantified (*n* = 16). All dots represent values for “Shoot length, cm” for individual plantlets, while red dots represent these values for plantlets that were chosen for pictures in A) and C). Box plots: red vertical line is median; box limits are lower and upper quartiles; whiskers are highest and lowest data points. Pairwise Wilcoxon Rank Sum Test with the p-value correction for multiple testing using Bonferroni adjustment method was used to calculate significant differences between groups.

### Selection of MV resistance in rice *lat5* calli

Transgenic and non-transgenic genome editing approaches in rice rely on callus induction, selection, and regeneration of shoots from transformed calli [9; 24]. To test whether the MV selection can be used during callus transformation to identify primary transformants, WT and *lat5* seeds were sown on callus-inducing medium supplemented either with solvent (0 μM MV), 0.1 μM MV or 1 μM MV. WT and *lat5* seeds germinated normally on all media. Six weeks after seeding *lat5* calli were formed on all tested media, while WT calli developed only on the callus-inducing media containing solvent and 0.1 μM MV (Fig 5). No WT calli were formed on the medium containing 1 μM MV. These data indicate that loss-of-function mutations in *LAT5* can be used as a selectable marker for primary transformants/editing events during callus formation stages either after transgenic or transgene-free genome editing.

**Fig 5.**
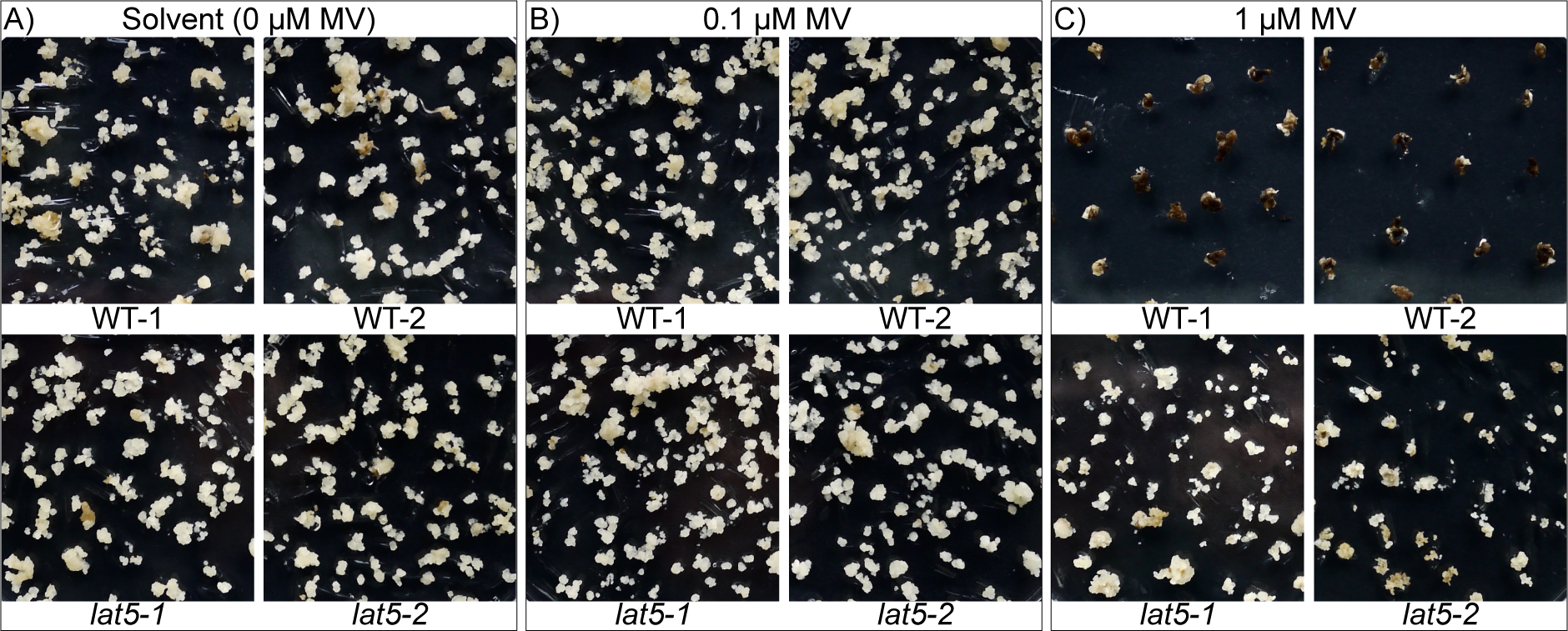
MV does not inhibit callus formation on *lat5* mature embryos. WT and loss-of-function *lat5* mature embryos were germinated on callus-inducing medium supplemented with either A) the solvent (0 μM MV), B) 0.1 μM MV or C) 1 μM MV. WT-1 and WT-2 are two independent lines of *O. sativa* cv. Kitaake harvested in 2022 and 2023, respectively. *lat5-1* and *lat5-2* are two independent lines homozygous for respective loss-of-function mutations in *LAT5* (Fig 2 and Table 1).

## Discussion

Transgene-based editing with CRISPR-Cas9 is presently a popular technique for introducing targeted mutations in model species and crops. Identification of transformant events relies on the introduction of selectable markers, typically genes encoding proteins that mediate resistance to antibiotics or herbicides. A selectable marker that can be obtained by editing, i.e., based on loss of function of an endogenous gene, would be advantageous for transgene-dependent as well as transgene-free genome editing approaches. Here, we developed a recessive marker that is caused by loss of function of an endogenous rice gene, i.e., *LAT5*/*PUT3*/*PAR1*, to confer resistance to the phytotoxin methylviologen (MV) as a method of selection. Note that, others used the LAT homolog *PAR1* from *Arabidopsis* to establish a similar selection system [14]. Also in parallel, Lyu *et al*. (2022) demonstrated that disruption of three rice polyamine uptake transporter genes, namely *LAT1*/*PUT1*, *LAT5*/*PUT3*/*PAR1*, and *LAT7*/*PUT2*, confers resistance to MV [29]. Our data indicate that disruption of a single polyamine uptake transporter gene in rice, i.e., *LAT5*/*PUT3*/*PAR1*, is sufficient to provide resistance to MV (Fig 3C, Fig 4C and Fig 5C). RNA-seq experiments show FPKM (Fragments Per Kilobase of transcript per Million mapped reads) values of 25 in both leaves and roots and >100 in calli for *LAT5*/*PUT3*/*PAR1* [35](S4A). By comparison, *LAT1*/*PUT1* had substantially lower mRNA levels in leaves roots and calli (S4B). *LAT7*/*PUT2* was also low in leaves and roots, but had FPKM values of about 24 in calli (S4C). The low base levels of the other two genes, namely *LAT1*/*PUT1* and *LAT7*/*PUT2*, in calli and leaves may explain why clear resistance to MV was only observed for *LAT5*/*PUT3*/*PAR1* (Fig 3C, Fig 4C and Fig 5C); however alternative hypotheses are also conceivable, e.g., different substrate specificity, lower transport capacity or different subcellular localization.

Because disruption of *LAT5*/*PUT3*/*PAR1* provides effective MV resistance (Fig 3C, Fig 4C and Fig 5C), gRNAs that target regions that encode transmembrane helices can be used for selecting transformants in transgene-based and transgene-free genome editing experiments. We demonstrate MV-based selection in photosynthetically active rice tissues, e.g., germinating seeds (Fig 3C) and young seedlings (Fig 4C). MV resistance of green tissues is consistent with MV being known to transfer electrons from photosystem I to molecular oxygen, which leads to formation of cytotoxic reactive oxygen species (ROS) and photodestruction of chlorophyll [25,37]. Notably, MV-based selection was also successfully obtained for non-photosynthetically active rice calli (Fig 5). Callus was grown photoautotrophically in a medium supplemented with 3% of sucrose. The high sucrose levels in the media prevented cytotoxic symptoms on the WT callus cells grown on the medium containing 0.1 μM MV [38] (Fig 5B), while at 1 μM MV callus induction from WT mature embryos was inhibited effectively (Fig 5C). Our results are in line with data from Zer et al. (1993)[39], who demonstrated that high MV concentrations inhibited growth of *Phaseolus vulgaris* cells during cultivation in darkness, likely by reducing DNA synthesis, and inhibiting the activity of enzymes involved in cellular defense against ROS. The deleterious effects of MV on non-photosynthetic cells was attributed to iron [39].

Since 1 μM MV blocked callus induction from WT mature embryos (Fig 5C), MV-based selection of *lat5*/*put3*/*par1* calli directly after transformation or transfection is possible. In the case of *Arabidopsis*, *PAR1* null alleles generated by CRISPR-Cas9 were shown to confer resistance to 1 µM and 10µM MV [14]. The authors stated that *heritable transgene-free mutations at target loci were identified in the T1 generation*. Apparently, although the inheritance of *par1* was considered to be recessive at 10µM MV, intermediate inheritance was be observed at lower 1 µM MV concentrations. In our study sequential reduction in MV concentrations enabled efficient and tight selection on 20-200-fold lower MV concentrations, i.e., 0.05 µM and 0.1 µM in rice.

Identification of lines that do not carry a transgene is an essential prerequisite for the classification of edited lines to be treated equivalent to under regulations [40]. One of the challenges is however that *Agrobacterium*-mediated transformation can lead to partial insertion of T-DNA copies [12]. Another challenge is the inadvertent insertion of vector backbone fragments into the genome of transformed plants [41,42]. Genotyping of hornless bulls generated with the help of genome editing identified the unintended insertion of plasmid sequences into the genome [43]. Thus, without further analyses, it is not possible to conclude that hygromycin-sensitive MV-resistant plants are transgene-free. Transgene-free editing based on transfection with CRISPR-enzyme-gRNA complexes or TALENs will be the new frontier to overcome the issues caused by *Agrobacterium*-mediated transformation.

Altogether we demonstrate that generation of loss-of-function mutations by editing in the single *OsLAT5/OsPUT3/OsPAR1* locus is sufficient to obtain resistance to MV and this selectable marker could be deployed for transgene-dependent and transgene-free genome editing approaches. Notably, we suggest that this approach can be easily adapted to generate orthogonal selection in other plants including Arabidopsis and crops.

## Acknowledgments

We would like to dedicate this article to Dr. Kazuo Shinozaki, whose pioneering work on methylviologen/polyamine transporters inspired this approach. We thank Dr. Michael Wudick for discussion and suggestions on experiments. This work was made possible by funding from the Bill and Melinda Gates Foundation to Heinrich Heine University, Düsseldorf, with a subawards to University of Missouri, Deutsche Forschungsgemeinschaft (DFG, German Research Foundation) under Germanýs Excellence Strategy – EXC-2048/1 – project ID 390686111 (CEPLAS), and an Alexander von Humboldt Professorship (WF). We also thank the Program for Promoting the Enhancement of Research Universities as young researcher units for the advancement of new and undeveloped fields at Nagoya University (MN). Note that in the course of this study two other groups developed analogous but different concepts

## Data availability

All data supporting the results are available in the main text or supplementary materials. The raw data will be deposited at DRYAD before publication. Materials will be made available under Material Transfer Agreement (MTA).

## Contributions

MN, BY and WF conceived of the study and supervised the project. Z.Z., B.L., and B.Y. assembled CRISPR constructs and developed single *OsLAT* knockout lines; K.S. conceived and designed experiments, analyzed the data and prepared figures; K.S., Z.Z, B.L., M.N., V.S.-L, E.P.I.L., B.Y., and W.B.F. wrote the manuscript.

## References

1. Tabassum J, Ahmad S, Hussain B, Mawia AM, Zeb A, Ju L. Applications and potential of genome-editing systems in rice improvement: current and future perspectives. Agronomy. 2021;11: 1359. doi:10.3390/agronomy11071359

2. Yang B. Grand challenges in genome editing in plants. Front Genome Ed. 2020;2. Available: https://www.frontiersin.org/articles/10.3389/fgeed.2020.00002

3. Eom J-S, Luo D, Atienza-Grande G, Yang J, Ji C, Luu VT, et al. Diagnostic kit for rice blight resistance. Nat Biotechnol. 2019;37: 1372–1379.

4. Oliva R, Ji C, Atienza-Grande G, Huguet-Tapia JC, Pérez-Quintéro A, Li T, et al. Broad-spectrum resistance to bacterial blight in rice using genome-editing. Nat Biotechnol. 2019;37: 1344–1350.

5. Schepler-Luu V, Sciallano C, Stiebner M, Ji C, Boulard G, Diallo A, et al. Genome editing of an African elite rice variety confers resistance against endemic and emerging *Xanthomonas oryzae pv. oryzae* strains. Weigel D, editor. eLife. 2023;12: e84864. doi:10.7554/eLife.84864

6. Gupta A, Liu B, Chen Q-J, Yang B. High-efficiency prime editing enables new strategies for broad-spectrum resistance to bacterial blight of rice. Plant Biotechnol J. 2023;21: 1454–1464. doi:10.1111/pbi.14049

7. Macovei A, Sevilla NR, Cantos C, Jonson GB, Slamet-Loedin I, Čermák T, et al. Novel alleles of rice eIF4G generated by CRISPR/Cas9-targeted mutagenesis confer resistance to Rice tungro spherical virus. Plant Biotechnol J. 2018;16: 1918–1927. doi:10.1111/pbi.12927

8. Wang F, Wang C, Liu P, Lei C, Hao W, Gao Y, et al. Enhanced rice blast resistance by CRISPR/Cas9-targeted Mutagenesis of the ERF transcription factor gene OsERF922. PloS One. 2016;11: e0154027. doi:10.1371/journal.pone.0154027

9. Arra Y, Auguy F, Stiebner M, Chéron S, Wudick MM, Miras M, et al. Rice Yellow Mottle Virus resistance by genome editing of the *Oryza sativa* L. *ssp. japonica* nucleoporin gene *OsCPR5.1* but not *OsCPR5.2*. bioRxiv; 2023. p. 2023.01.13.523077. doi:10.1101/2023.01.13.523077

10. Sharma A, Chouhan A, Bhatt T, Kaur A, Minhas AP. Selectable Markers to Marker-Free Selection in Rice. Mol Biotechnol. 2022;64: 841–851. doi:10.1007/s12033-022-00460-w

11. Shimada TL, Shimada T, Hara-Nishimura I. A rapid and non-destructive screenable marker, FAST, for identifying transformed seeds of *Arabidopsis thaliana*. Plant J. 2010;61: 519–528. doi:10.1111/j.1365-313X.2009.04060.x

12. Sun L, Ge Y, Sparks JA, Robinson ZT, Cheng X, Wen J, et al. TDNAscan: A software to identify complete and truncated T-DNA insertions. Front Genet. 2019;10:685. doi:doi: 10.3389/fgene.2019.00685

13. Gu X, Liu L, Zhang H. Transgene-free genome editing in plants. Front Genome Ed. 2021;3. Available: https://www.frontiersin.org/articles/10.3389/fgeed.2021.805317

14. Kong X, Pan W, Zhang T, Liu L, Zhang H. A simple and efficient strategy to produce transgene-free gene edited plants in one generation using paraquat resistant 1 as a selection marker. Front Plant Sci. 2023;13. Available: https://www.frontiersin.org/articles/10.3389/fpls.2022.1051991

15. Aliaga-Franco N, Zhang C, Presa S, Srivastava AK, Granell A, Alabadí D, et al. Identification of Transgene-Free CRISPR-Edited Plants of Rice, Tomato, and Arabidopsis by Monitoring DsRED Fluorescence in Dry Seeds. Front Plant Sci. 2019;10. doi:10.3389/fpls.2019.01150

16. Kim Y-A, Moon H, Park C-J. CRISPR/Cas9-targeted mutagenesis of *Os8N3* in rice to confer resistance to *Xanthomonas oryzae pv. oryzae*. Rice N Y N. 2019;12: 67. doi:10.1186/s12284-019-0325-7

17. Toda E, Koiso N, Takebayashi A, Ichikawa M, Kiba T, Osakabe K, et al. An efficient DNA- and selectable-marker-free genome-editing system using zygotes in rice. Nat Plants. 2019;5: 363. doi:10.1038/s41477-019-0386-z

18. Andersson M, Turesson H, Olsson N, Fält A-S, Ohlsson P, Gonzalez MN, et al. Genome editing in potato via CRISPR-Cas9 ribonucleoprotein delivery. Physiol Plant. 2018;164: 378–384. doi:10.1111/ppl.12731

19. Liang Z, Chen K, Li T, Zhang Y, Wang Y, Zhao Q, et al. Efficient DNA-free genome editing of bread wheat using CRISPR/Cas9 ribonucleoprotein complexes. Nat Commun. 2017;8. doi:10.1038/ncomms14261

20. Park S-C, Park S, Jeong YJ, Lee SB, Pyun JW, Kim S, et al. DNA-free mutagenesis of *GIGANTEA* in *Brassica oleracea var. capitata* using CRISPR/Cas9 ribonucleoprotein complexes. Plant Biotechnol Rep. 2019;13: 483–489. doi:10.1007/s11816-019-00585-6

21. Sant’Ana RRA, Caprestano CA, Nodari RO, Agapito-Tenfen SZ. PEG-delivered CRISPR-Cas9 ribonucleoproteins system for gene-editing screening of maize protoplasts. Genes. 2020;11: 1029. doi:10.3390/genes11091029

22. Fujita M, Fujita Y, Iuchi S, Yamada K, Kobayashi Y, Urano K, et al. Natural variation in a polyamine transporter determines paraquat tolerance in Arabidopsis. Proc Natl Acad Sci U S A. 2012;109: 6343–6347. doi:10.1073/pnas.1121406109

23. Fujita M, Shinozaki K. Identification of polyamine transporters in plants: paraquat transport provides crucial clues. Plant Cell Physiol. 2014;55: 855–861. doi:10.1093/pcp/pcu032

24. Shen Y, Ruan Q, Chai H, Yuan Y, Yang W, Chen J, et al. The Arabidopsis polyamine transporter LHR1/PUT3 modulates heat responsive gene expression by enhancing mRNA stability. Plant J. 2016;88: 1006–1021. doi:10.1111/tpj.13310

25. Li J, Mu J, Bai J, Fu F, Zou T, An F, et al. Paraquat Resistant1, a Golgi-localized putative transporter protein, is involved in intracellular transport of paraquat. Plant Physiol. 2013;162: 470–483. doi:10.1104/pp.113.213892

26. Mulangi V, Phuntumart V, Aouida M, Ramotar D, Morris P. Functional analysis of OsPUT1, a rice polyamine uptake transporter. Planta. 2012;235: 1–11. doi:10.1007/s00425-011-1486-9

27. Xi J, Xu P, Xiang C-B. Loss of AtPDR11, a plasma membrane-localized ABC transporter, confers paraquat tolerance in Arabidopsis thaliana. Plant J. 2012;69: 782–791. doi:10.1111/j.1365-313X.2011.04830.x

28. Nazish T, Huang Y-J, Zhang J, Xia J-Q, Alfatih A, Luo C, et al. Understanding paraquat resistance mechanisms in Arabidopsis thaliana to facilitate the development of paraquat-resistant crops. Plant Commun. 2022;3: 100321. doi:10.1016/j.xplc.2022.100321

29. Lyu Y-S, Cao L-M, Huang W-Q, Liu J-X, Lu H-P. Disruption of three polyamine uptake transporter genes in rice by CRISPR/Cas9 gene editing confers tolerance to herbicide paraquat. aBIOTECH. 2022;3: 140–145. doi:10.1007/s42994-022-00075-4

30. Luu VT, Stiebner M, Maldonado PE, Valdés S, Marín D, Delgado G, et al. Efficient Agrobacterium-mediated transformation of the elite–indica rice variety Komboka. Bio-Protoc. 2020;10: e3739–e3739.

31. Zhou H, Liu B, Weeks DP, Spalding MH, Yang B. Large chromosomal deletions and heritable small genetic changes induced by CRISPR/Cas9 in rice. Nucleic Acids Res. 2014;42: 10903– 10914. doi:10.1093/nar/gku806

32. Wang P, Khoshravesh R, Karki S, Tapia R, Balahadia CP, Bandyopadhyay A, et al. Re-creation of a key step in the evolutionary switch from C3 to C4 leaf anatomy. Curr Biol CB. 2017;27: 3278–3287.e6. doi:10.1016/j.cub.2017.09.040

33. Murray MG, Thompson WF. Rapid isolation of high molecular weight plant DNA. Nucleic Acids Res. 1980;8: 4321–4325.

34. Zhao H, Ma H, Yu L, Wang X, Zhao J. Genome-wide survey and expression analysis of amino acid transporter gene family in rice (*Oryza sativa* L.). PLoS ONE. 2012;7: e49210. doi:10.1371/journal.pone.0049210

35. Xia L, Zou D, Sang J, Xu X, Yin H, Li M, et al. Rice Expression Database (RED): An integrated RNA-Seq-derived gene expression database for rice. J Genet Genomics Yi Chuan Xue Bao. 2017;44: 235–241. doi:10.1016/j.jgg.2017.05.003

36. Mulangi GRV. Characterization of polyamine transporters from rice and *Arabidopsis* - ProQuest. PhD thesis, Bowling Green State University. 2011. Available: https://www.proquest.com/openview/089ccfea528706874366a9630d420aa1/1?cbl=18750&pq-

37. Farrington JA, Ebert M, Land EJ, Fletcher K. Bipyridylium quaternary salts and related compounds. V. Pulse radiolysis studies of the reaction of paraquat radical with oxygen. Implications for the mode of action of bipyridyl herbicides. Biochim Biophys Acta BBA - Bioenerg. 1973;314: 372–381. doi:10.1016/0005-2728(73)90121-7

38. Moreland DE, Hilton JL. Actions on photosynthetic systems. In: Audus LJ, editor. Herbicides: physiology, biochemistry, ecology. London, New York: Academic Press; 1976.

39. Zer H, Chevion M, Goldberg I. Effect of paraquat on dark-grown *Phaseolus vulgaris* cells. Weed Sci. 1993;41: 528–533.

40. Buchholzer M, Frommer WB. An increasing number of countries regulate genome editing in crops. New Phytol. 2023;237: 12–15. doi:10.1111/nph.18333

41. De Buck S, De Wilde C, Van Montagu M, Depicker A. T-DNA vector backbone sequences are frequently integrated into the genome of transgenic plants obtained by Agrobacterium-mediated transformation. Mol Breed. 2000;6: 459–468. doi:10.1023/A:1026575524345

42. Kononov ME, Bassuner B, Gelvin SB. Integration of T-DNA binary vector “backbone” sequences into the tobacco genome: evidence for multiple complex patterns of integration. Plant J. 1997;11: 945–57.

43. Young AE, Mansour TA, McNabb BR, Owen JR, Trott JF, Brown CT, et al. Genomic and phenotypic analyses of six offspring of a genome-edited hornless bull. Nat Biotechnol. 2020;38: 225–232. doi:10.1038/s41587-019-0266-0

