## Supplementary Data for "Loss-of-function mutation in the polyamine transporter gene *OsLAT5* as a selectable marker for genome editing"

**S1 Table. Oligonucleotides used for assembly of CRISPR constructs and for genotyping of putative transformants.**

| # | Oligo name | Sequence 5'>3' | Purpose |
|---|---|---|---|
| 1 | gLAT5-F1 | gcagGAAGAGCGTAAGGGAGGCCA | Assembly of CRISPR construct for *OsLAT5* knockout |
| 2 | gLAT5-R1 | aaacTGGCCTCCCTTACGCTCTTC | |
| 3 | gLAT5-F2 | gcagTTCTGCTGTTTGCACTCATA | |
| 4 | gLAT5-R2 | aaacTATGAGTGCAAACAGCAGAA | |
| 5 | gLAT1-F1 | gcagCGACTCGTACCAGCTGCT | Assembly of CRISPR construct for *OsLAT1* knockout |
| 6 | gLAT1-R1 | aaacAGCAGCTGGTACGAGTCG | |
| 7 | gLAT1-F2 | gcagGATGAAGGCGACGAACTCG | |
| 8 | gLAT1-R2 | aaacCGAGTTCGTCGCCTTCATC | |
| 9 | gLAT7-F1 | gcagTGCTCTGGTCCCTCCCCG | Assembly of CRISPR construct for *OsLAT7* knockout |
| 10 | gLAT7-R1 | aaacCGGGGAGGGACCAGAGCA | |
| 11 | gLAT7-F2 | gcagACGAGGAACGCCGCGGC | |
| 12 | gLAT7-R2 | aaacGCCGCGGCGTTCCTCGT | |
| 13 | LAT5-F1 | GTCAACTTAAAACTCGGATATG | PCR-amplification of *OsLAT5* for genotyping CRISPR plants |
| 14 | LAT5-R1 | ACCCAGCCAACTATTGTCAA | |
| 15 | LAT1-F1 | CTACTGGGATTCGATCAGC | PCR-amplification of *OsLAT1* for genotyping CRISPR plants |
| 16 | LAT1-R1 | AGATCCGGGTTAACGGAGAA | |
| 17 | LAT7-F1 | CTCGTCGCGCTCATCTTCTA | PCR-amplification of *OsLAT7* for genotyping CRISPR plants |
| 18 | LAT7-R1 | CTCGCCTTGTCCCAGTAGTT | |

**S2 Appendix. Alignment of coding sequences of *AtPAR1*, *OsLAT1*, *OsLAT5*, and *OsLAT7*.** Alignments are made using MUSCLE (Multiple Sequence Comparison by Log- Expectation) algorithm. Nucleotides that are conserved in 75% of sequences are shaded in BLACK using pyBoxshade program. Sequences encoding transmembrane domains of AtPAR1 are colored in **blue** according to UniProt database (Q9C6S5 · PHSB_ARATH). Two guide RNAs were used for knock out of each rice gene. Selected guide RNAs are colored in **red**, while PAM sites are colored in **green**.

```
           ....:...10....:...20....:...30....:...40....:...50....:...60
AtPAR1     ATGCA--GAAGCGGAGAAT---CATCACTGTGAA--------------------CCCCTC
OsLAT1     ATGGCGGACACCGGCGGACGGCCGGAGGTGTCGCTGGCCACGGTCCGGTCGCCGGGCCAC
OsLAT5     ATGAC--GAACGCATGGAT---C-TCGCCGTCGGTGGTCGCCCTCTGCCCCTCTCCCCTC
OsLAT7     ATGACCGGAGCCTGCGAGG---CGGCGCCGGC----------------------------

           ....:...70....:...80....:...90....:..100....:..110....:..120
AtPAR1     C-----------------------------------------------------------
OsLAT1     CCGGCAGCTTCTACGACG------------------------------------------
OsLAT5     CCCTCATCTCGCCTCCCGGGTTCTGTCCTCTCCTGCTGGCCGGACAGCCGCGGCATCCGG
OsLAT7     ------------------------------------------------------------

           ....:..130....:..140....:..150....:..160....:..170....:..180
AtPAR1     -----------------------------------GCGTCTATTGAGATGAGTCA--G--
OsLAT1     -----------------------------------GCAGCGGCGGCGGCGGATCTCGGCC
OsLAT5     CGCGGGGCTGGCGAGGGGACAGCCGGACAGACGCTGCGGCCGGCGAGGGGATTCACTG--
OsLAT7     -----------------------------------GCGGCGGCGGGG-------------

           ....:..190....:..200....:..210....:..220....:..230....:..240
AtPAR1     -------TACGAGAATAATGAAGTTC--------------CTTACTCAA-----------
OsLAT1     ACGCTGACACCGGGCAAGAGAAGCCCACCGTCGAGAGCGCCCAACC--------------
OsLAT5     -------TAGAGAAATTAAGGAACACAGCAATAACACGAGCTAACTCAGCCTGTCTTCCA
OsLAT7     ------------------------------------------------------------

           ....:..250....:..260....:..270....:..280....:..290....:..300
AtPAR1     ----------GTGTCG-------------GTGCTG-ATGAGGTTC--CATCATCACC---
OsLAT1     --GGCGAACGGTGCCGCTCCGATGGGCGAGTGCGGCACGGAGTACAGGGGCCTCCCCGAC
OsLAT5     ATGGAGGATTGTGTTG-------------GT-----ATCAAGTACAGCAGTGTCAATGAG
OsLAT7     ------------------------------------------------------------

           ....:..310....:..320....:..330....:..340....:..350....:..360
AtPAR1     --------GCCTAAAGCTACCGATAAGATT-------CGGAAAGTTTCCATGTTGCCACT
OsLAT1     GGCG----ACGCCGGCGGGCCAATGCCGTCGTCGGCACGCACGGTTTCGATGATCCCGCT
OsLAT5     GGCGAAGAGCGTAAGGGAGGCCATGGCGTC-------CCAAAGGTTTCCATCATCCCACT
OsLAT7     -------------------------------------CTGACGG------TGCTCCCCCT

           ....:..370....:..380....:..390....:..400....:..410....:..420
AtPAR1     TGTTTTCCTCATATTCTATGAAGTCTCAGGAGGTCCTTTTGGTGTAGAGGATAGTGTGAA
OsLAT1     CATCTTCCTCATCTTCTACGAGGTGTCCGGCGGGCCGTTCGGGATCGAGGACAGCGTGGG
OsLAT5     CATTTTCCTCATATTCTATGAAGTTTCTGGGGGTCCGTTTGGGATTGAGGATAGTGTCAA
OsLAT7     CGTCGCGCTCATCTTCTACGACGTGTCGGGGGGCCCCTTCGGCATCGAGGACTCGGTCCG

           ....:..430....:..440....:..450....:..460....:..470....:..480
AtPAR1     TGC---AGCAGGACCATTATTAGCTCTTCTAGGGTTTGTGATCTTTCCTTTCATTTGGAG
OsLAT1     CGC---GGCCGGGCCGCTGCTCGCCATCATCGGCTTCTTGGTCCTCCCCGTCATCTGGAG
OsLAT5     GGC---TGCTGGCCCACTCCTAGCAATTGCTGGATTTCTGCTGTTTGCACTCATATGGAG
OsLAT7     CGCCGGCGGCGGCGCGCTCCTCCCGATCCTGGGGTTCCTCGTCCTCCCCGTGCTCTGGTC
```

```
             ....:..490....:..500....:..510....:..520....:..530....:..540
AtPAR1       TATCCCTGAGGCTTTGATTACTGCGGAGATGGGAACAATGTATCCCGAGAATGGTGGTTA
OsLAT1       CATCCCGGAGGCGCTGATCACGGCGGAGCTGGGCGCCATGTTCCCGGAGAACGGCGGGTA
OsLAT5       TGTCCCGGAAGCCCTGATTACTGCAGAGATGGGCACTATGTTTCCTGAGAATGGTGGTTA
OsLAT7       CCTCCCCGAGGCGCTCGTCACCGCCGAGCTCGCCTCCGCGTTCCCCACCAACGCCGGCTA

             ....:..550....:..560....:..570....:..580....:..590....:..600
AtPAR1       TGTTGTGTGGGTTTCCTCTGCTTTGGACCTTTTTGGGGGTTTCAGCAAGGTTGGATGAA
OsLAT1       CGTCGTGTGGGTGGCGTCGGCGCTCGGCCCGTACTGGGGGTTCCAGCAAGGGTGGATGAA
OsLAT5       CGTCGTCTGGGTCTCTTCAGCCCTTGGCCATTCTGGGGTTTTCAGCAAGGCTGGCAAA
OsLAT7       CGTCGCCTGGGTCTCCGCCGCGTTCGGCCCCGCCGCGGCGTTCCTCGTCGGGTTCTCCAA

             ....:..610....:..620....:..630....:..640....:..650....:..660
AtPAR1       ATGGCTTAGTGGTGTTATTGATAACGCTTTGTATCCTGTTCTGTTTCTTGACTATTTGAA
OsLAT1       GTGGTTGAGCGGCGTCATCGACAACGCGCTCTACCCCGTCCTCTTCTTGGACTACCTCAA
OsLAT5       GTGGCTGAGTGGTGTCATAGATAATGCTCTCTATCCAGTCCTCTTCCTCGACTATGTTAA
OsLAT7       GTGGGCGTCGGGACGCTCGACAACGCGCTCTACCCGGTGCTCTTCCTCGACTACCTCCG

             ....:..670....:..680....:..690....:..700....:..710....:..720
AtPAR1       GTCTGGA-----GTCCCGGCTTTAGGTAGCGGTTTACCGAGAGTTGCATCGATCTTGGTG
OsLAT1       GTCCGGC-----GTCCCGGCGCTCGGCGGAGGCGCGCCGAGGGCGTTCGCCGTCGTCGGC
OsLAT5       GTCCAGC-----ATTCCAGCTCTTGGAGGTGGTCTCCCAAGGACCTTGGCGGTGCTTATC
OsLAT7       CTCCGGCGGGGGCTCGTGCTCTCCCCGCCGGCCCGCT-----CCCTCGCCGTGCTCGCG

             ....:..730....:..740....:..750....:..760....:..770....:..780
AtPAR1       CTGACTATTTTGTTGACGTATCTGAACTACAGGGGACTAACTATAGTTGGTTGGGTTGCT
OsLAT1       CTGACGGCCGTGCTGACATTGCTGAATTACCGGGGGCTCACCGTCGTCGGATGGGTGGCG
OsLAT5       CTCACAGTTGCACTTACTTACATGAACTACAGAGGGTTGACAATAGTTGGCTGGGTGGCA
OsLAT7       CTCACCGCCGCGCTCACCTACCTCAACTTCCGGGGGCTCCACCTCGTCGGCCTCTCCGCG

             ....:..790....:..800....:..810....:..820....:..830....:..840
AtPAR1       GTGCTTATGGGAGTTTTCTCCATCCTTCCGTTTGCTGTGATGGGTTTGATTTCGATTCCG
OsLAT1       ATCTGCCTTGGCGTCTTCTCCCTCCTCCCTTTTTTCGTCATGGGGCTCATCGCGCTCCCC
OsLAT5       GTCTTTCTTGGCGTGTTCTCTCTACTCCCGTTTTTTGTTATGGGATTAATAGCTATTCCC
OsLAT7       CTGGCGCTCACCGCGTTCTCGCTCTCCCCGTTCGTCGCGCTCGCCGTGCTCGCCGCCCCC

             ....:..850....:..860....:..870....:..880....:..890....:..900
AtPAR1       CAGCTAGAGCCTTCGAGATGGCTTGTGATGGACTTAGGGAATGTGAACTGGAATTTGTAT
OsLAT1       AAGCTCCGGCCGGCGAGGTGGCTCGTGATCGACCTCCACAACGTCGATTGGAATCTGTAC
OsLAT5       CGAATCGAACCCTCAAGATGGCTTGAAATGGACTTGGGGAATGTGAATTGGGGTTTATAT
OsLAT7       AAGATCCGCCCGTCGCGGTGGCTCGCCGTGAACGTGGCCGCCGTTGAGCCGCGCGCCTAC

             ....:..910....:..920....:..930....:..940....:..950....:..960
AtPAR1       CTTAACACGCTTTTCTGGAATCTAAACTATTGGGACTCGATTAGTACACTAGCTGGTGAA
OsLAT1       CTGAACACTCTGTTCTGGAACCTCAACTACTGGGATTCGATCAGCACGTTGGCCGGCGAG
OsLAT5       CTAAACACACTGTTTTGGAACCTCAATTATTGGGACTCAATCAGTACCCTTGCTGGAGAG
OsLAT7       TTCAACTCCATGTTCTGGAACCTCAACTACTGGGACAAGGCGAGCACGCTTGCCGGCGAG

             ....:..970....:..980....:..990....:.1000....:.1010....:.1020
AtPAR1       GTTGAAAACCCCAACCATACACTTCCAAAGGCTTTGTTTTATGGTGTGATCTTAGTTGCT
OsLAT1       GTGAAGAATCCCGGCAAGACGCTGCCCAAGGCGCTGTTCTACGCGGTCATCTTCGTGGTG
OsLAT5       GTTGAGAATCCAAAGAGAACACTCCCAAGGGCACTTTCTTATGCTCTAGTTTTAGTGGTG
OsLAT7       GTGGAGGAGCCGAGGAAGACGTTCCCGAAGGCGGTGTTCGGCGCGGTGGGGCTCGTCGTG
```

```
        ....:.1030....:.1040....:.1050....:.1060....:.1070....:.1080
AtPAR1  TGTTCTTACATCTTCCCTCTTTTGGCTGGAATCGGGGCAATCCCGCTAGAGC---GTGAG
OsLAT1  GTCGCCTACCTGTACCCTCTCCTCGCCGGGACGGGAGCCGTGCCGCTGGACA---GGGGG
OsLAT5  GGGGGATACCTCTACCCTCTGATCACCTGTACAGCAGCAGTTCCAGTTGTTC---GGGAG
OsLAT7  GGCGCGTACCTCATCCCGCTCCTCGCCGGGACGGGCGCGCTGCCGTCGGAGACGGCGGGG

        ....:.1090....:.1100....:.1110....:.1120....:.1130....:.1140
AtPAR1  AAATGGACAGATGGGTATTTTTCTGATGTGGCCAAAGCTCTTGGTGGAGCGTGGTTGAGA
OsLAT1  CAGTGGACAGACGGCTACTTCGCGGACATCGCGAAGCTGCTCGGCGGCGCGTGGCTGATG
OsLAT5  TTCTGGACGGATGGATATTTCTCAGACGTTGCGAGAATTCTTGGTGGTTTCTGGTTGCAC
OsLAT7  GAGTGGACGGACGGGTTCTTCTCCGTGGTCGGCGACCGGATCGGCGGGCCGTGGCTGCGC

        ....:.1150....:.1160....:.1170....:.1180....:.1190....:.1200
AtPAR1  TGGTGGGTTCAAGCGGCTGCAGCTACATCAAACATGGGAATGTTTATAGCCGAGATGAGT
OsLAT1  TGGTGGGTGCAGTCGGCGGCGGCGCTGTCGAACATGGGCATGTTCGTGGCGGAGATGAGC
OsLAT5  TCGTGGCTTCAAGCAGCTGCTGCACTGTCCAACATGGGCAATTTCGTAACTGAAATGAGC
OsLAT7  GTGTGGATCCAGGCCGCCGCGGCCATGTCCAACATGGGGCTCTTCGAGGCCGAGATGAGC

        ....:.1210....:.1220....:.1230....:.1240....:.1250....:.1260
AtPAR1  AGTGACTCTTTTCAGCTTCTCGGAATGGCTGAGCGCGGTATGCTTCCCGAGTTCTTTGCC
OsLAT1  AGCGACTCGTACCAGCTGCTGGGCATGGCGGAGCGGGGCATGCTCCCGTCCTTCTTCGCG
OsLAT5  AGTGATTCTTACCAGCTTCTCGGGATGGCTGAGCGTGGAATGCTTCCAGAGTTTTTCGCC
OsLAT7  GGCGACTCGTTCCAGCTCCTCGGCATGGCGGAGATGGGCATGATCCCGGCGATCTTCGCG

        ....:.1270....:.1280....:.1290....:.1300....:.1310....:.1320
AtPAR1  AAAAGGTCACGTTACGGGACACCTTTACTAGGGATTCTGTTTTCAGCGTCAGGTGTTGTT
OsLAT1  GCGCGGTCGCGGTACGGCACGCCGCTGGCGGGCATCCTCTTCTCGGCCTCCGGCGTGCTG
OsLAT5  AAGAGATCTCGCTATGGAACCCCACTTATTGGCATCATGTTCTCCGCGTTTGGTGTGGTC
OsLAT7  CGCAGGTCGCGCCACGGCACGCCGACGTACAGCATCCTCTGCTCGGCCACCGGCGTCGTC

        ....:.1330....:.1340....:.1350....:.1360....:.1370....:.1380
AtPAR1  CTCTTGTCTTGGCTAAGCTTTCAAGAGATTGTAGCTGCAGAGAACTTACTCTACTGCGTT
OsLAT1  CTGCTCTCGATGATGAGCTTCCAGGAGATCGTGGCGGCCGAGAACTTCCTCTACTGCTTC
OsLAT5  CTGCTGTCCTGGATGAGCTTCCAGGAGATCATCGCTGCGGAGAACTACCTGTACTGCTTC
OsLAT7  ATCCTCTCCTTCATGAGCTTCCAGGAGATCGTCGAGTTCCTCAACTTCCTCTACGGCCTC

        ....:.1390....:.1400....:.1410....:.1420....:.1430....:.1440
AtPAR1  GGTATGATCTTAGAGTTTATAGCGTTTGTTAGAATGAGAATGAAACACCCTGCTGCATCA
OsLAT1  GGCATGCTCCTCGAGTTCGTCGCCTTCATCCTGCACCGGGTGAGGCGCCCCGACGCGGCG
OsLAT5  GGTATGATCCTGGAATTCATCGCCTTCATCAAGCTGAGGGTGGTCCACCCAAACGCCTCC
OsLAT7  GGGATGCTCGCCGTGTTCGCCGCCTTCGTCAAGCTCCGCGTCAAGGACCCCGACCTCCCC

        ....:.1450....:.1460....:.1470....:.1480....:.1490....:.1500
AtPAR1  AGACCGTACAAGATACCTATTGGAACCACAGGTTCGATCCTCATGTGTATTCCTCCAACC
OsLAT1  CGCCCATACAGGGTGCCGCTGGGCACAGCCGGGTGCGTGGCGATGCTGGTGCCGCCGACG
OsLAT5  CGACCTTACAAGATCCCACTGGGCACCATCGGCGCTGTCCTGATGATCATCCCACCTACC
OsLAT7  CGCCCGTACCGGATCCCCGTCGGCGCCGCGGGCGCCGCCGCCATGTGCGTCCCGCCCGTC

        ....:.1510....:.1520....:.1530....:.1540....:.1550....:.1560
AtPAR1  ATACTGATCTGTGCTGTTGTAGCACTCTCGTCTCTTAAAGTAGCTGCAGTGAGCATTGTG
OsLAT1  GCGCTGATCGCCGTGGTGCTCGCGCTGTCCACGCTGAAGGTGGCGGTGGTGAGCCTCGGC
OsLAT5  ATTCTGATCGTCGTGGTGATGATGCTCGCGTCCTTCAAGGTGATGGTGGTCAGCATCATG
OsLAT7  GTCCTCATCACCACCGTCATGTGCCTCGCCTCCGCCAGGACGCTCGTCGTCAGCGCCGCC
```

```
       ....:.1570....:.1580....:.1590....:.1600....:.1610....:.1620
AtPAR1 ATGATGATCATTGGTTTCCTGATACATCCTTTACTGAACCATATGGATCGAAAGAGATGG
OsLAT1 GCGGTGGCCATGGGGCTCGTGCTGCAGCCGGCGCTGAGGTTCGTGGAGAAGAAGCGGTGG
OsLAT5 GCAATGCTGGTTGGGTTCGTGCTGCAGCCGGCTCTGGTGTACGTGGAGAAGAGACGGTGG
OsLAT7 GTGGCCGTCGCCGGCGTCGCCATGTACTACGGCGTCGAGCACATGAAGGCCACCGGCTGC

       ....:.1630....:.1640....:.1650....:.1660....:.1670....:.1680
AtPAR1 GTCAAATTCTC--------------------------------------------------
OsLAT1 CTGAGGTTCTC--------------------------------------------------
OsLAT5 CTGAAGTTCTC--------------------------------------------------
OsLAT7 GTCGAGTTCTTGACGCCGGTGCCGCCTGACAGCCTCCGTGGATCATCATCATCATCATCC

       ....:.1690....:.1700....:.1710....:.1720....:.1730....:.1740
AtPAR1 -----------------------CATCAGTTCTGACTTACCGGA----------------
OsLAT1 -----------------------CGTTAACCCGGATCTCCCGGAGATCGGCGTGATTCGC
OsLAT5 -----------------------CATAAGCGCAGAACTGCCAGA----------------
OsLAT7 TCATCGGCAGCGTCCGACAACGGCGGCGACGACGACGTCGAGGACGTCTGCGCCCTCCTC

       ....:.1750....:.1760....:.1770....:.1780....:.1790....:..
AtPAR1 CCTTCAACA---GCAGACTCGGGAATACGAAGAAACTCTA---ATA-----CGTTAA
OsLAT1 CCGCCCGC-------CGCGCCGGACGAGCCGTTGGTTCCG--------------TAG
OsLAT5 TTTGCCGTACTCGAACGTTGAGGAAGACAGCACAATCCCACTTGTG-----TGCTGA
OsLAT7 CTCGCCGCCGGCGAGCACGCCGGAGAAGGCGTCAGTGTCAGCAAGGAGAATTATTAG
```

**S3 Appendix. Alignment of AtPAR1, OsLAT1, OsLAT5, and OsLAT7 protein sequences.** Alignments are made using Clustal Omega algorithm. Amino acids that are conserved in 75% of sequences are shaded in BLACK using pyBoxshade program. Transmembrane domains of AtPAR1 are colored in **blue** according to UniProt database (Q9C6S5 · PHSB_ARATH).

```
        ....:...10...:...20...:...30...:...40...:...50...:...60
AtPAR1 ---------------------------------------------------------MQK
OsLAT1 MADTGGRPEV------SLATVRSPGHPAASTTAA--AAADLGHADTGQEKP-----TVE-
OsLAT5 MTNAWISPSVVALCPSPLPSSRLPGSVLSCWPDSRGIRRGAGEGTAGQTLRPARGFTVEK
OsLAT7 ------------------------------------------------------------

        ....:...70...:...80...:...90...:..100...:..110...:..120
AtPAR1 RR--IITVNPSASIEMSQYENNEVPYSSVGADEVPSSPPKATDKIRKVSMLPLVFLIFYE
OsLAT1 -----SAQPANGAAPM---GECGTEYRGLPDGDAGG---PMPSSARTVSMIPLIFLIFYE
OsLAT5 LRNTAITRANSACLPM--EDCVGIKYSSVNEG----EERKGGHGVPKVSIIPLIFLIFYE
OsLAT7 ---------------------------------MTGACEAAPARRRGLTVLPLVALIFYD

        ....:..130...:..140...:..150...:..160...:..170...:..180
AtPAR1 VSGGPFGVEDSVNAA-GPLLALLGFVIFPFIWSIPEALITAEMGTMYPENGGYVVWVSSA
OsLAT1 VSGGPFGIEDSVGAA-GPLLAIIGFLVLPVIWSIPEALITAELGAMFPENGGYVVWVASA
OsLAT5 VSGGPFGIEDSVKAA-GPLLAIAGFLLFALIWSVPEALITAEMGTMFPENGGYVVWVSSA
OsLAT7 VSGGPFGIEDSVRAGGALLPILGFLVLPVLWSLPEALVTAELASAFPTNAGYVAWVSAA

        ....:..190...:..200...:..210...:..220...:..230...:..240
AtPAR1 LGPFWGFQQGWMKWLSGVIDNALYPVLFLDYLKSGVPALGSGLPRVASILVLTILLTYLN
OsLAT1 LGPYWGFQQGWMKWLSGVIDNALYPVLFLDYLKSGVPALGGAPRAFAVVGLTAVLTLLN
OsLAT5 LGPFWGFQQGWAKWLSGVIDNALYPVLFLDYVKSSIPALGGGLPRTLAVLILTVALTYMN
OsLAT7 FGPAAAFLVGFSKWASGTLDNALYPVLFLDYLRSGGGLVLSPPARSLAVLALTAALTYLN

        ....:..250...:..260...:..270...:..280...:..290...:..300
AtPAR1 YRGLTIVGWVAVLMGVFSILPFAVMGLISIPQLEPSRWLVMDLGNVNWNLYLNTLFWNLN
OsLAT1 YRGLTVVGWVAICLGVFSLLPFFVMGLIALPKLRPARWLVIDLHNVDWNLYLNTLFWNLN
OsLAT5 YRGLTIVGWVAVFLGVFSLLPFFVMGLIAIPRIEPSRWLEMDLGNVNWGLYLNTLFWNLN
OsLAT7 FRGLHLVGLSALALTAFSLSPFVALAVLAAPKIRPSRWLAVNVAAVEPRAYFNSMFWNLN

        ....:..310...:..320...:..330...:..340...:..350...:..360
AtPAR1 YWDSISTLAGEVENPNHTLPKALFYGVILVACSYIFPLLAGIGAIPLE-REKWTDGYFSD
OsLAT1 YWDSISTLAGEVKNPGKTLPKALFYAVIFVVVAYLYPLLAGTGAVPLD-RGQWTDGYFAD
OsLAT5 YWDSISTLAGEVENPKRTLPRALSYALVLVVGGYLYPLITCTAAVPVV-REFWTDGYFSD
OsLAT7 YWDKASTLAGEVEEPRKTFPKAVFGAVGLVVGAYLIPLLAGTGALPSETAGEWTDGFFSV

        ....:..370...:..380...:..390...:..400...:..410...:..420
AtPAR1 VAKALGGAWLRWWVQAAAATSNMGMFIAEMSSDSFQLLGMAERGMLPEFFAKRSRYGTPL
OsLAT1 IAKLLGGAWLMWWVQSAAALSNMGMFVAEMSSDSYQLLGMAERGMLPSFFAARSRYGTPL
OsLAT5 VARILGGFWLHSWLQAAAALSNMGNFVTEMSSDSYQLLGMAERGMLPEFFAKRSRYGTPL
OsLAT7 VGDRIGGPWLRVWIQAAAAMSNMGLFEAEMSGDSFQLLGMAEMGMIPAIFARRSRHGTPT

        ....:..430...:..440...:..450...:..460...:..470...:..480
AtPAR1 LGILFSASGVVLLSWLSFQEIVAAENLLYCVGMILEFIAFVRMRMKHPAASRPYKIPIGT
OsLAT1 AGILFSASGVLLLSMMSFQEIVAAENFLYCFGMLLEFVAFILHRVRRPDAARPYRVPLGT
OsLAT5 IGIMFSAFGVVLLSWMSFQEIIAAENYLYCFGMILEFIAFIKLRVVHPNASRPYKIPLGT
OsLAT7 YSILCSATGVVILSFMSFQEIVEFLNFLYGLGMLAVFAAFVKLRVKDPDLPRPYRIPVGA
```

```
        ....:..490....:..500....:..510....:..520....:..530....:..540
AtPAR1  TGSILMCIPPTILICAVVALSSSLKVAAVSIVMMIIGFLIHPLLNHMDRKRWVKFSISSDL
OsLAT1  AGCVAMLVPPTALIAVVLALSTLKVAVVSLGAVAMGLVLQPALRFVEKKRWLRFSVNPDL
OsLAT5  IGAVLMIIPPTILIVVVMMLASFKVMVVSIMAMLVGFVLQPALVYVEKRRWLKFSISAEL
OsLAT7  AGAAAMCVPPVVLITTVMCLASARTLVVSAAVAVAGVAMYYGVEHMKATGCVEFLTPVPP

        ....:..550....:..560....:..570....:..580....:..590
AtPAR1  PDLQQQ----------TREYEETLIR-----------------------*
OsLAT1  PEIGVI----------RPPAAPDEPLVP---------------------*
OsLAT5  PDLPYS----------NVEEDSTIPLVC---------------------*
OsLAT7  DSLRGSSSSSSSSAASDNGGDDDVEDVCALLLAAGEHAGEGVSVSKENY*
```

**S4 Appendix. Organ-specific mRNA level for (A) *OsLAT5*/*OsPUT3*/*OsPAR1*, (B) *OsLAT1*/*OsPUT1* and (C) *OsLAT7*/*OsPUT2*** [31]. FPKM: Fragments Per Kilobase of transcript per Million mapped reads.

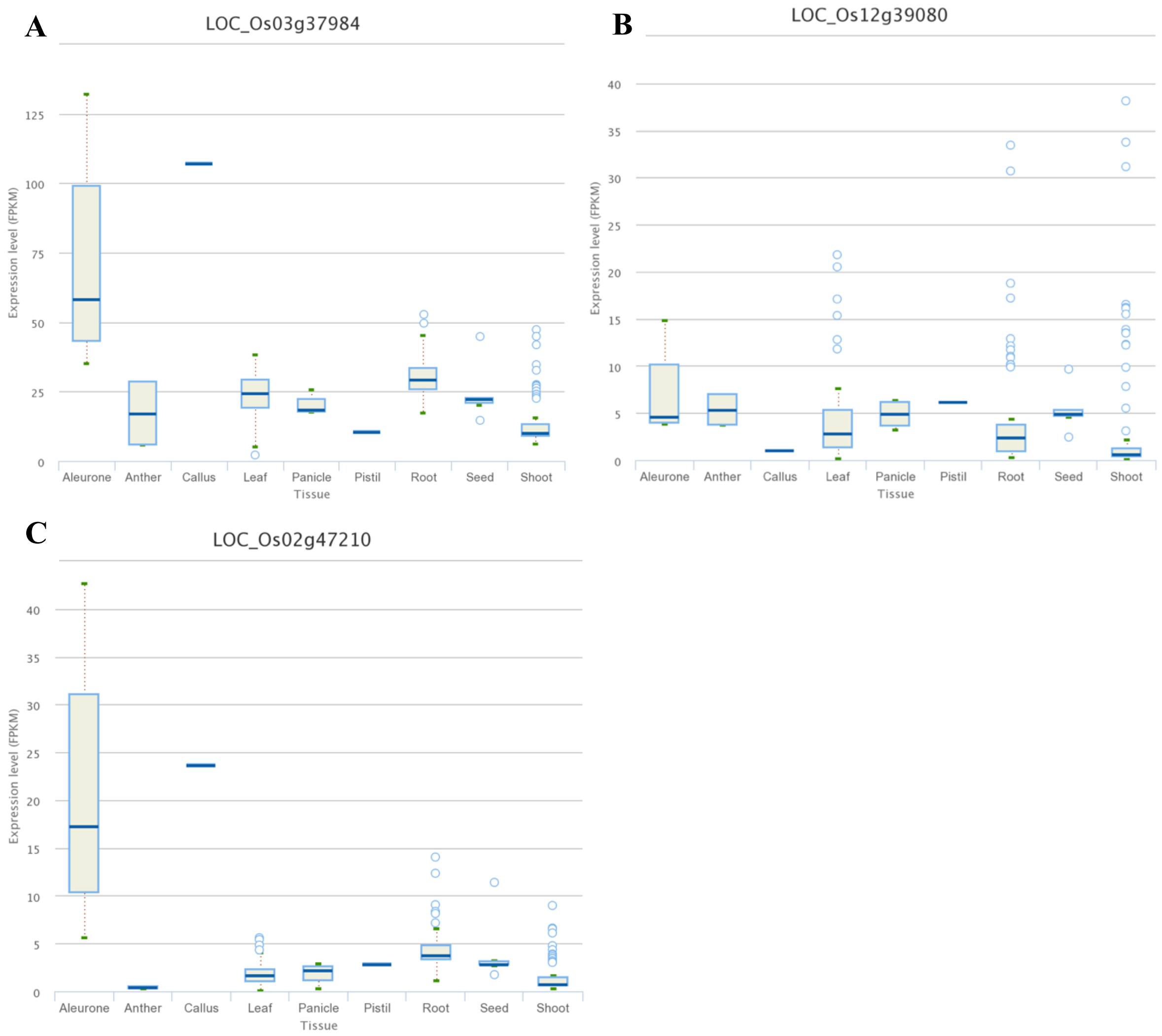

**S5 Appendix. Alignments of coding sequences (CDS) of rice *LAT1*, *LAT5*, *LAT7* wild-type and the mutant alleles used in this study.** Alignments are made using MUSCLE (Multiple Sequence Comparison by Log- Expectation) algorithm. Non-mutated nucleotides that are shaded in BLACK using pyBoxshade program. Deleted nucleotides in mutant alleles are represented with – symbol. Inserted nucleotides in mutant alleles are non-shaded. Two guide RNAs were used for knock out of each rice gene. Selected guide RNAs are colored in **red**, while PAM sites are colored in **green**.

### *LAT1/PUT1* (Os02g0700500; LOC_Os02g47210)

```
           ....:...10...:...20...:...30...:...40...:...50...:...60
LAT1       ATGGCGGACACCGGCGGACGGCCGGAGGTGTCGCTGGCCACGGTCCGGTCGCCGGGCCAC
lat1-1     ATGGCGGACACCGGCGGACGGCCGGAGGTGTCGCTGGCCACGGTCCGGTCGCCGGGCCAC
lat1-2     ATGGCGGACACCGGCGGACGGCCGGAGGTGTCGCTGGCCACGGTCCGGTCGCCGGGCCAC

           ....:...70...:...80...:...90...:..100...:..110...:..120
LAT1       CCGGCAGCTTCTACGACGGCAGCGGCGGCGGCGGATCTCGGCCACGCTGACACCGGGCAA
lat1-1     CCGGCAGCTTCTACGACGGCAGCGGCGGCGGCGGATCTCGGCCACGCTGACACCGGGCAA
lat1-2     CCGGCAGCTTCTACGACGGCAGCGGCGGCGGCGGATCTCGGCCACGCTGACACCGGGCAA

           ....:..130...:..140...:..150...:..160...:..170...:..180
LAT1       GAGAAGCCCACCGTCGAGAGCGCCCAACCGGCGAACGGTGCCGCTCCGATGGGCGAGTGC
lat1-1     GAGAAGCCCACCGTCGAGAGCGCCCAACCGGCGAACGGTGCCGCTCCGATGGGCGAGTGC
lat1-2     GAGAAGCCCACCGTCGAGAGCGCCCAACCGGCGAACGGTGCCGCTCCGATGGGCGAGTGC

           ....:..190...:..200...:..210...:..220...:..230...:..240
LAT1       GGCACGGAGTACAGGGGCCTCCCCGACGGCGACGCCGGCGGGCCAATGCCGTCGTCGGCA
lat1-1     GGCACGGAGTACAGGGGCCTCCCCGACGGCGACGCCGGCGGGCCAATGCCGTCGTCGGCA
lat1-2     GGCACGGAGTACAGGGGCCTCCCCGACGGCGACGCCGGCGGGCCAATGCCGTCGTCGGCA

           ....:..250...:..260...:..270...:..280...:..290...:..300
LAT1       CGCACGGTTTCGATGATCCCGCTCATCTTCCTCATCTTCTACGAGGTGTCCGGCGGGCCG
lat1-1     CGCACGGTTTCGATGATCCCGCTCATCTTCCTCATCTTCTACGAGGTGTCCGGCGGGCCG
lat1-2     CGCACGGTTTCGATGATCCCGCTCATCTTCCTCATCTTCTACGAGGTGTCCGGCGGGCCG

           ....:..310...:..320...:..330...:..340...:..350...:..360
LAT1       TTCGGGATCGAGGACAGCGTGGGCGCGGCCGGGCCGCTGCTCGCCATCATCGGCTTCTTG
lat1-1     TTCGGGATCGAGGACAGCGTGGGCGCGGCCGGGCCGCTGCTCGCCATCATCGGCTTCTTG
lat1-2     TTCGGGATCGAGGACAGCGTGGGCGCGGCCGGGCCGCTGCTCGCCATCATCGGCTTCTTG

           ....:..370...:..380...:..390...:..400...:..410...:..420
LAT1       GTCCTCCCCGTCATCTGGAGCATCCCGGAGGCGCTGATCACGGCGGAGCTGGGCGCCATG
lat1-1     GTCCTCCCCGTCATCTGGAGCATCCCGGAGGCGCTGATCACGGCGGAGCTGGGCGCCATG
lat1-2     GTCCTCCCCGTCATCTGGAGCATCCCGGAGGCGCTGATCACGGCGGAGCTGGGCGCCATG

           ....:..430...:..440...:..450...:..460...:..470...:..480
LAT1       TTCCCGGAGAACGGCGGGTACGTCGTGTGGGTGGCGTCGGCGCTCGGCCCGTACTGGGGG
lat1-1     TTCCCGGAGAACGGCGGGTACGTCGTGTGGGTGGCGTCGGCGCTCGGCCCGTACTGGGGG
lat1-2     TTCCCGGAGAACGGCGGGTACGTCGTGTGGGTGGCGTCGGCGCTCGGCCCGTACTGGGGG

           ....:..490...:..500...:..510...:..520...:..530...:..540
LAT1       TTCCAGCAAGGGTGGATGAAGTGGTTGAGCGGCGTCATCGACAACGCGCTCTACCCCGTC
lat1-1     TTCCAGCAAGGGTGGATGAAGTGGTTGAGCGGCGTCATCGACAACGCGCTCTACCCCGTC
lat1-2     TTCCAGCAAGGGTGGATGAAGTGGTTGAGCGGCGTCATCGACAACGCGCTCTACCCCGTC


           ....:..550...:..560...:..570...:..580...:..590...:..600
```

```
LAT1   CTCTTCTTGGACTACCTCAAGTCCGGCGTCCCGGCGCTCGGCGGAGGCGCGCCGAGGGCG
lat1-1 CTCTTCTTGGACTACCTCAAGTCCGGCGTCCCGGCGCTCGGCGGAGGCGCGCCGAGGGCG
lat1-2 CTCTTCTTGGACTACCTCAAGTCCGGCGTCCCGGCGCTCGGCGGAGGCGCGCCGAGGGCG

       ....:..610....:..620....:..630....:..640....:..650....:..660
LAT1   TTCGCCGTCGTCGGCCTGACGGCCGTGCTGACATTGCTGAATTACCGGGGGCTCACCGTC
lat1-1 TTCGCCGTCGTCGGCCTGACGGCCGTGCTGACATTGCTGAATTACCGGGGGCTCACCGTC
lat1-2 TTCGCCGTCGTCGGCCTGACGGCCGTGCTGACATTGCTGAATTACCGGGGGCTCACCGTC

       ....:..670....:..680....:..690....:..700....:..710....:..720
LAT1   GTCGGATGGGTGGCGATCTGCCTTGGCGTCTTCTCCCTCCTCCCTTTTTCGTCATGGGG
lat1-1 GTCGGATGGGTGGCGATCTGCCTTGGCGTCTTCTCCCTCCTCCCTTTTTCGTCATGGGG
lat1-2 GTCGGATGGGTGGCGATCTGCCTTGGCGTCTTCTCCCTCCTCCCTTTTTCGTCATGGGG

       ....:..730....:..740....:..750....:..760....:..770....:..780
LAT1   CTCATCGCGCTCCCCAAGCTCCGGCCGGCGAGGTGGCTCGTGATCGACCTCCACAACGTC
lat1-1 CTCATCGCGCTCCCCAAGCTCCGGCCGGCGAGGTGGCTCGTGATCGACCTCCACAACGTC
lat1-2 CTCATCGCGCTCCCCAAGCTCCGGCCGGCGAGGTGGCTCGTGATCGACCTCCACAACGTC

       ....:..790....:..800....:..810....:..820....:..830....:..840
LAT1   GATTGGAATCTGTACCTGAACACTCTGTTCTGGAACCTCAACTACTGGGATTCGATCAGC
lat1-1 GATTGGAATCTGTACCTGAACACTCTGTTCTGGAACCTCAACTACTGGGATTCGATCAGC
lat1-2 GATTGGAATCTGTACCTGAACACTCTGTTCTGGAACCTCAACTACTGGGATTCGATCAGC

       ....:..850....:..860....:..870....:..880....:..890....:..900
LAT1   ACGTTGGCCGGCGAGGTGAAGAATCCCGGCAAGACGCTGCCCAAGGCGCTGTTCTACGCG
lat1-1 ACGTTGGCCGGCGAGGTGAAGAATCCCGGCAAGACGCTGCCCAAGGCGCTGTTCTACGCG
lat1-2 ACGTTGGCCGGCGAGGTGAAGAATCCCGGCAAGACGCTGCCCAAGGCGCTGTTCTACGCG

       ....:..910....:..920....:..930....:..940....:..950....:..960
LAT1   GTCATCTTCGTGGTGGTCGCCTACCTGTACCCTCTCCTCGCCGGGACGGAGCCGTGCCG
lat1-1 GTCATCTTCGTGGTGGTCGCCTACCTGTACCCTCTCCTCGCCGGGACGGAGCCGTGCCG
lat1-2 GTCATCTTCGTGGTGGTCGCCTACCTGTACCCTCTCCTCGCCGGGACGGAGCCGTGCCG

       ....:..970....:..980....:..990....:.1000....:.1010....:.1020
LAT1   CTGGACAGGGGGCAGTGGACAGACGGCTACTTCGCGGACATCGCGAAGCTGCTCGGCGGC
lat1-1 CTGGACAGGGGGCAGTGGACAGACGGCTACTTCGCGGACATCGCGAAGCTGCTCGGCGGC
lat1-2 CTGGACAGGGGGCAGTGGACAGACGGCTACTTCGCGGACATCGCGAAGCTGCTCGGCGGC

       ....:.1030....:.1040....:.1050....:.1060....:.1070....:.1080
LAT1   GCGTGGCTGATGTGGTGGGTGCAGTCGGCGGCGGCGCTGTCGAACATGGGCATGTTCGTG
lat1-1 GCGTGGCTGATGTGGTGGGTGCAGTCGGCGGCGGCGCTGTCGAACATGGGCATGTTCGTG
lat1-2 GCGTGGCTGATGTGGTGGGTGCAGTCGGCGGCGGCGCTGTCGAACATGGGCATGTTCGTG

       ....:.1090....:.1100....:.1110....:.1120....:.1130....:.1140
LAT1   GCGGAGATGAGCAGCGACTCGTACCAGC-TGCTGGGCATGGCGGAGCGGGGCATGCTCCC
lat1-1 GCGGAGATGAGCAGCGACTCGTA------TGCTGGGCATGGCGGAGCGGGGCATGCTCCC
lat1-2 GCGGAGATGAGCAGCGACTCGTACCAGCTTGCTGGGCATGGCGGAGCGGGGCATGCTCCC

       ....:.1150....:.1160....:.1170....:.1180....:.1190....:.1200
LAT1   GTCCTTCTTCGCGGCGCGGTCGCGGTACGGCACGCCGCTGGCGGGCATCCTCTTCTCGGC
lat1-1 GTCCTTCTTCGCGGCGCGGTCGCGGTACGGCACGCCGCTGGCGGGCATCCTCTTCTCGGC
lat1-2 GTCCTTCTTCGCGGCGCGGTCGCGGTACGGCACGCCGCTGGCGGGCATCCTCTTCTCGGC
```

```
        ....:.1210....:.1220....:.1230....:.1240....:.1250....:.1260
LAT1    CTCCGGCGTGCTGCTGCTCTCGATGATGAGCTTCCAGGAGATCGTGGCGGCCGAGAACTT
lat1-1  CTCCGGCGTGCTGCTGCTCTCGATGATGAGCTTCCAGGAGATCGTGGCGGCCGAGAACTT
lat1-2  CTCCGGCGTGCTGCTGCTCTCGATGATGAGCTTCCAGGAGATCGTGGCGGCCGAGAACTT

        ....:.1270....:.1280....:.1290....:.1300....:.1310....:.1320
LAT1    CCTCTACTGCTTCGGCATGCTCCTCGAGTTCGTCGCCTTCATCCTGCACCGGGTGAGGCG
lat1-1  CCTCTACTGCTTCGGCATGCTCCTCGAGTTCGTCGCCTTCATCCTGCACCGGGTGAGGCG
lat1-2  CCTCTACTGCTTCGGCATGCTCCTCGAGTTCGTCGCCTTCATCCTGCACCGGGTGAGGCG

        ....:.1330....:.1340....:.1350....:.1360....:.1370....:.1380
LAT1    CCCCGACGCGGCGCGCCCATACAGGGTGCCGCTGGGCACAGCCGGGTGCGTGGCGATGCT
lat1-1  CCCCGACGCGGCGCGCCCATACAGGGTGCCGCTGGGCACAGCCGGGTGCGTGGCGATGCT
lat1-2  CCCCGACGCGGCGCGCCCATACAGGGTGCCGCTGGGCACAGCCGGGTGCGTGGCGATGCT

        ....:.1390....:.1400....:.1410....:.1420....:.1430....:.1440
LAT1    GGTGCCGCCGACGGCGCTGATCGCCGTGGTGCTCGCGCTGTCCACGCTGAAGGTGGCGGT
lat1-1  GGTGCCGCCGACGGCGCTGATCGCCGTGGTGCTCGCGCTGTCCACGCTGAAGGTGGCGGT
lat1-2  GGTGCCGCCGACGGCGCTGATCGCCGTGGTGCTCGCGCTGTCCACGCTGAAGGTGGCGGT

        ....:.1450....:.1460....:.1470....:.1480....:.1490....:.1500
LAT1    GGTGAGCCTCGGCGCGGTGGCCATGGGGCTCGTGCTGCAGCCGGCGCTGAGGTTCGTGGA
lat1-1  GGTGAGCCTCGGCGCGGTGGCCATGGGGCTCGTGCTGCAGCCGGCGCTGAGGTTCGTGGA
lat1-2  GGTGAGCCTCGGCGCGGTGGCCATGGGGCTCGTGCTGCAGCCGGCGCTGAGGTTCGTGGA

        ....:.1510....:.1520....:.1530....:.1540....:.1550....:.1560
LAT1    GAAGAAGCGGTGGCTGAGGTTCTCCGTTAACCCGGATCTCCCGGAGATCGGCGTGATTCG
lat1-1  GAAGAAGCGGTGGCTGAGGTTCTCCGTTAACCCGGATCTCCCGGAGATCGGCGTGATTCG
lat1-2  GAAGAAGCGGTGGCTGAGGTTCTCCGTTAACCCGGATCTCCCGGAGATCGGCGTGATTCG

        ....:.1570....:.1580....:.1590....:..
LAT1    CCCGCCCGCCGCGCCGGACGAGCCGTTGGTTCCGTAG
lat1-1  CCCGCCCGCCGCGCCGGACGAGCCGTTGGTTCCGTAG
lat1-2  CCCGCCCGCCGCGCCGGACGAGCCGTTGGTTCCGTAG
```

***LAT5/PUT3/PAR1*** **(Os03g0576900; LOC_Os03g37984)**

```
        ....:...10....:...20....:...30....:...40....:...50....:...60
LAT5    ATGACGAACGCATGGATCTCGCCGTCGGTGGTCGCCCTCTGCCCCTCTCCCCTCCCCTCA
lat5-1  ATGACGAACGCATGGATCTCGCCGTCGGTGGTCGCCCTCTGCCCCTCTCCCCTCCCCTCA
lat5-2  ATGACGAACGCATGGATCTCGCCGTCGGTGGTCGCCCTCTGCCCCTCTCCCCTCCCCTCA

        ....:...70....:...80....:...90....:..100....:..110....:..120
LAT5    TCTCGCCTCCCGGGTTCTGTCCTCTCCTGCTGGCCGGACAGCCGCGGCATCCGGCGCGGG
lat5-1  TCTCGCCTCCCGGGTTCTGTCCTCTCCTGCTGGCCGGACAGCCGCGGCATCCGGCGCGGG
lat5-2  TCTCGCCTCCCGGGTTCTGTCCTCTCCTGCTGGCCGGACAGCCGCGGCATCCGGCGCGGG

        ....:..130....:..140....:..150....:..160....:..170....:..180
LAT5    GCTGGCGAGGGGACAGCCGGACAGACGCTGCGGCCGGCGAGGGGATTCACTGTAGAGAAA
lat5-1  GCTGGCGAGGGGACAGCCGGACAGACGCTGCGGCCGGCGAGGGGATTCACTGTAGAGAAA
lat5-2  GCTGGCGAGGGGACAGCCGGACAGACGCTGCGGCCGGCGAGGGGATTCACTGTAGAGAAA
```

```
        ....:..190....:..200....:..210....:..220....:..230....:..240
LAT5    TTAAGGAACACAGCAATAACACGAGCTAACTCAGCCTGTCTTCCAATGGAGGATTGTGTT
lat5-1  TTAAGGAACACAGCAATAACACGAGCTAACTCAGCCTGTCTTCCAATGGAGGATTGTGTT
lat5-2  TTAAGGAACACAGCAATAACACGAGCTAACTCAGCCTGTCTTCCAATGGAGGATTGTGTT

        ....:..250....:..260....:..270....:..280....:..290....:..300
LAT5    GGTATCAAGTACAGCAGTGTCAATGAGGGCGAAGAGCGTAAGGGAGGCCATGGCGTCCCA
lat5-1  GGTATCAAGTACAGCAGTGTCAATGAGGGCGAAGAGCGTAAGGGA-GCCATGGCGTCCCA
lat5-2  GGTATCAAGTACAGCAGTGTCAATGAGGGCGAAGAGCGTAAGGGA-------GCGTCCCA

        ....:..310....:..320....:..330....:..340....:..350....:..360
LAT5    AAGGTTTCCATCATCCCACTCATTTTCCTCATATTCTATGAAGTTTCTGGGGGTCCGTTT
lat5-1  AAGGTTTCCATCATCCCACTCATTTTCCTCATATTCTATGAAGTTTCTGGGGGTCCGTTT
lat5-2  AAGGTTTCCATCATCCCACTCATTTTCCTCATATTCTATGAAGTTTCTGGGGGTCCGTTT

        ....:..370....:..380....:..390....:..400....:..410....:..420
LAT5    GGGATTGAGGATAGTGTCAAGGCTGCTGGCCCACTCCTAGCAATTGCTGGATTTCTGCTG
lat5-1  GGGATTGAGGATAGTGTCAAGGCTGCTGGCCCACTCCTAGCA----------------TG
lat5-2  GGGATTGAGGATAGTGTCAAGGCTGCTGGCCCACTCCTAGCAATTGCTGGATTTCTGCTG

        ....:..430....:..440....:..450....:..460....:..470....:..480
LAT5    TTTGCACTCATATGGAGTGTCCCGGAAGCCCTGATTACTGCAGAGATGGCACTATGTTT
lat5-1  ATCACAATT-TCTGGAGTGTCCCGGAAGCCCTGATTACTGCAGAGATGGCACTATGTTT
lat5-2  TTTGCACTCATATGGAGTGTCCCGGAAGCCCTGATTACTGCAGAGATGGCACTATGTTT

        ....:..490....:..500....:..510....:..520....:..530....:..540
LAT5    CCTGAGAATGGTGGTTACGTCGTCTGGGTCTCTTCAGCCCTTGGCCATTCTGGGGTTTT
lat5-1  CCTGAGAATGGTGGTTACGTCGTCTGGGTCTCTTCAGCCCTTGGCCATTCTGGGGTTTT
lat5-2  CCTGAGAATGGTGGTTACGTCGTCTGGGTCTCTTCAGCCCTTGGCCATTCTGGGGTTTT

        ....:..550....:..560....:..570....:..580....:..590....:..600
LAT5    CAGCAAGGCTGGGCAAAGTGGCTGAGTGGTGTCATAGATAATGCTCTCTATCCAGTCCTC
lat5-1  CAGCAAGGCTGGGCAAAGTGGCTGAGTGGTGTCATAGATAATGCTCTCTATCCAGTCCTC
lat5-2  CAGCAAGGCTGGGCAAAGTGGCTGAGTGGTGTCATAGATAATGCTCTCTATCCAGTCCTC

        ....:..610....:..620....:..630....:..640....:..650....:..660
LAT5    TTCCTCGACTATGTTAAGTCCAGCATTCCAGCTCTTGGAGGTGGTCTCCCAAGGACCTTG
lat5-1  TTCCTCGACTATGTTAAGTCCAGCATTCCAGCTCTTGGAGGTGGTCTCCCAAGGACCTTG
lat5-2  TTCCTCGACTATGTTAAGTCCAGCATTCCAGCTCTTGGAGGTGGTCTCCCAAGGACCTTG

        ....:..670....:..680....:..690....:..700....:..710....:..720
LAT5    GCGGTGCTTATCCTCACAGTTGCACTTACTTACATGAACTACAGAGGGTTGACAATAGTT
lat5-1  GCGGTGCTTATCCTCACAGTTGCACTTACTTACATGAACTACAGAGGGTTGACAATAGTT
lat5-2  GCGGTGCTTATCCTCACAGTTGCACTTACTTACATGAACTACAGAGGGTTGACAATAGTT

        ....:..730....:..740....:..750....:..760....:..770....:..780
LAT5    GGCTGGGTGGCAGTCTTTCTTGGCGTGTTCTCTCTACTCCCGTTTTTTGTTATGGGATTA
lat5-1  GGCTGGGTGGCAGTCTTTCTTGGCGTGTTCTCTCTACTCCCGTTTTTTGTTATGGGATTA
lat5-2  GGCTGGGTGGCAGTCTTTCTTGGCGTGTTCTCTCTACTCCCGTTTTTTGTTATGGGATTA

        ....:..790....:..800....:..810....:..820....:..830....:..840
LAT5    ATAGCTATTCCCCGAATCGAACCCTCAAGATGGCTTGAAATGGACTTGGGGAATGTGAAT
lat5-1  ATAGCTATTCCCCGAATCGAACCCTCAAGATGGCTTGAAATGGACTTGGGGAATGTGAAT
lat5-2  ATAGCTATTCCCCGAATCGAACCCTCAAGATGGCTTGAAATGGACTTGGGGAATGTGAAT
```

```
        ....:..850....:..860....:..870....:..880....:..890....:..900
LAT5    TGGGGTTTATATCTAAACACACTGTTTTGGAACCTCAATTATTGGGACTCAATCAGTACC
lat5-1  TGGGGTTTATATCTAAACACACTGTTTTGGAACCTCAATTATTGGGACTCAATCAGTACC
lat5-2  TGGGGTTTATATCTAAACACACTGTTTTGGAACCTCAATTATTGGGACTCAATCAGTACC

        ....:..910....:..920....:..930....:..940....:..950....:..960
LAT5    CTTGCTGGAGAGGTTGAGAATCCAAAGAGAACACTCCCAAGGGCACTTTCTTATGCTCTA
lat5-1  CTTGCTGGAGAGGTTGAGAATCCAAAGAGAACACTCCCAAGGGCACTTTCTTATGCTCTA
lat5-2  CTTGCTGGAGAGGTTGAGAATCCAAAGAGAACACTCCCAAGGGCACTTTCTTATGCTCTA

        ....:..970....:..980....:..990....:.1000....:.1010....:.1020
LAT5    GTTTTAGTGGTGGGGGATACCTCTACCCTCTGATCACCTGTACAGCAGCAGTTCCAGTT
lat5-1  GTTTTAGTGGTGGGGGATACCTCTACCCTCTGATCACCTGTACAGCAGCAGTTCCAGTT
lat5-2  GTTTTAGTGGTGGGGGATACCTCTACCCTCTGATCACCTGTACAGCAGCAGTTCCAGTT

        ....:.1030....:.1040....:.1050....:.1060....:.1070....:.1080
LAT5    GTTCGGGAGTTCTGGACGGATGGATATTTCTCAGACGTTGCGAGAATTCTTGGTGGTTTC
lat5-1  GTTCGGGAGTTCTGGACGGATGGATATTTCTCAGACGTTGCGAGAATTCTTGGTGGTTTC
lat5-2  GTTCGGGAGTTCTGGACGGATGGATATTTCTCAGACGTTGCGAGAATTCTTGGTGGTTTC

        ....:.1090....:.1100....:.1110....:.1120....:.1130....:.1140
LAT5    TGGTTGCACTCGTGGCTTCAAGCAGCTGCTGCACTGTCCAACATGGGCAATTTCGTAACT
lat5-1  TGGTTGCACTCGTGGCTTCAAGCAGCTGCTGCACTGTCCAACATGGGCAATTTCGTAACT
lat5-2  TGGTTGCACTCGTGGCTTCAAGCAGCTGCTGCACTGTCCAACATGGGCAATTTCGTAACT

        ....:.1150....:.1160....:.1170....:.1180....:.1190....:.1200
LAT5    GAAATGAGCAGTGATTCTTACCAGCTTCTCGGGATGGCTGAGCGTGGAATGCTTCCAGAG
lat5-1  GAAATGAGCAGTGATTCTTACCAGCTTCTCGGGATGGCTGAGCGTGGAATGCTTCCAGAG
lat5-2  GAAATGAGCAGTGATTCTTACCAGCTTCTCGGGATGGCTGAGCGTGGAATGCTTCCAGAG

        ....:.1210....:.1220....:.1230....:.1240....:.1250....:.1260
LAT5    TTTTTCGCCAAGAGATCTCGCTATGGAACCCCACTTATTGGCATCATGTTCTCCGCGTTT
lat5-1  TTTTTCGCCAAGAGATCTCGCTATGGAACCCCACTTATTGGCATCATGTTCTCCGCGTTT
lat5-2  TTTTTCGCCAAGAGATCTCGCTATGGAACCCCACTTATTGGCATCATGTTCTCCGCGTTT

        ....:.1270....:.1280....:.1290....:.1300....:.1310....:.1320
LAT5    GGTGTGGTCCTGCTGTCCTGGATGAGCTTCCAGGAGATCATCGCTGCGGAGAACTACCTG
lat5-1  GGTGTGGTCCTGCTGTCCTGGATGAGCTTCCAGGAGATCATCGCTGCGGAGAACTACCTG
lat5-2  GGTGTGGTCCTGCTGTCCTGGATGAGCTTCCAGGAGATCATCGCTGCGGAGAACTACCTG

        ....:.1330....:.1340....:.1350....:.1360....:.1370....:.1380
LAT5    TACTGCTTCGGTATGATCCTGGAATTCATCGCCTTCATCAAGCTGAGGGTGGTCCACCCA
lat5-1  TACTGCTTCGGTATGATCCTGGAATTCATCGCCTTCATCAAGCTGAGGGTGGTCCACCCA
lat5-2  TACTGCTTCGGTATGATCCTGGAATTCATCGCCTTCATCAAGCTGAGGGTGGTCCACCCA

        ....:.1390....:.1400....:.1410....:.1420....:.1430....:.1440
LAT5    AACGCCTCCCGACCTTACAAGATCCCACTGGGCACCATCGGCGCTGTCCTGATGATCATC
lat5-1  AACGCCTCCCGACCTTACAAGATCCCACTGGGCACCATCGGCGCTGTCCTGATGATCATC
lat5-2  AACGCCTCCCGACCTTACAAGATCCCACTGGGCACCATCGGCGCTGTCCTGATGATCATC

        ....:.1450....:.1460....:.1470....:.1480....:.1490....:.1500
LAT5    CCACCTACCATTCTGATCGTCGTGGTGATGATGCTCGCGTCCTTCAAGGTGATGGTGGTC
lat5-1  CCACCTACCATTCTGATCGTCGTGGTGATGATGCTCGCGTCCTTCAAGGTGATGGTGGTC
lat5-2  CCACCTACCATTCTGATCGTCGTGGTGATGATGCTCGCGTCCTTCAAGGTGATGGTGGTC
```

```
        ....:.1510....:.1520....:.1530....:.1540....:.1550....:.1560
LAT5    AGCATCATGGCAATGCTGGTTGGGTTCGTGCTGCAGCCGGCTCTGGTGTACGTGGAGAAG
lat5-1  AGCATCATGGCAATGCTGGTTGGGTTCGTGCTGCAGCCGGCTCTGGTGTACGTGGAGAAG
lat5-2  AGCATCATGGCAATGCTGGTTGGGTTCGTGCTGCAGCCGGCTCTGGTGTACGTGGAGAAG

        ....:.1570....:.1580....:.1590....:.1600....:.1610....:.1620
LAT5    AGACGGTGGCTGAAGTTCTCCATAAGCGCAGAACTGCCAGATTTGCCGTACTCGAACGTT
lat5-1  AGACGGTGGCTGAAGTTCTCCATAAGCGCAGAACTGCCAGATTTGCCGTACTCGAACGTT
lat5-2  AGACGGTGGCTGAAGTTCTCCATAAGCGCAGAACTGCCAGATTTGCCGTACTCGAACGTT

        ....:.1630....:.1640....:.1650...
LAT5    GAGGAAGACAGCACAATCCCACTTGTGTGCTGA
lat5-1  GAGGAAGACAGCACAATCCCACTTGTGTGCTGA
lat5-2  GAGGAAGACAGCACAATCCCACTTGTGTGCTGA
```

***LAT7/PUT2*** **(Os12g0580400; LOC_Os12g39080)**

```
        ....:...10....:...20....:...30....:...40....:...50....:...60
LAT7    ATGACCGGAGCCTGCGAGGCGGCGCCGGCGCGGCGGCGGGGGCTGACGGTGCTCCCCCTC
lat7-1  ATGACCGGAGCCTGCGAGGCGGCGCCGGCGCGGCGGCGGGGGCTGACGGTGCTCCCCCTC
lat7-2  ATGACCGGAGCCTGCGAGGCGGCGCCGGCGCGGCGGCGGGGGCTGACGGTGCTCCCCCTC

        ....:...70....:...80....:...90....:..100....:..110....:..120
LAT7    GTCGCGCTCATCTTCTACGACGTGTCGGGGGGCCCCTTCGGCATCGAGGACTCGGTCCGC
lat7-1  GTCGCGCTCATCTTCTACGACGTGTCGGGGGGCCCCTTCGGCATCGAGGACTCGGTCCGC
lat7-2  GTCGCGCTCATCTTCTACGACGTGTCGGGGGGCCCCTTCGGCATCGAGGACTCGGTCCGC

        ....:..130....:..140....:..150....:..160....:..170....:..180
LAT7    GCCGGCGGCGGCGCGCTCCTCCCGATCCTGGGGTTCCTCGTCCTCCCCGTGCTCTGGTCC
lat7-1  GCCGGCGGCGGCGCGCTCCTCCCGATCCTGGGGTTCCTCGTCCTCCCCGTGCTCTGGTCC
lat7-2  GCCGGCGGCGGCGCGCTCCTCCCGATCCTGGGGTTCCTCGTCCTCCCCGTGCTCTGGTC-

        ....:..190....:..200....:..210....:..220....:..230....:..240
LAT7    CTCCCCGAGGCGCTCGTCACCGCCGAGCTCGCCTCCGCGTTCCCCACCAACGCCGGCTAC
lat7-1  CT-CCCGAGGCGCTCGTCACCGCCGAGCTCGCCTCCGCGTTCCCCACCAACGCCGGCTAC
lat7-2  ---CCCGAGGCGCTCGTCACCGCCGAGCTCGCCTCCGCGTTCCCCACCAACGCCGGCTAC

        ....:..250....:..260....:..270....:..280....:..290....:..300
LAT7    GTCGCCTGGGTCTCCGCCGCGTTCGGCCCCGCCGCGGCGTTCCTCGTCGGGTTCTCCAAG
lat7-1  GTCGCCTGGGTCTCCGCCGCGTTCGGCCCCGCCGCGGCGTTCCTCGTCGGGTTCTCCAAG
lat7-2  GTCGCCTGGGTCTCCGCCGCGTTCGGCCCCGCCGCGGCGTTCCTCGTCGGGTTCTCCAAG

        ....:..310....:..320....:..330....:..340....:..350....:..360
LAT7    TGGGCGTCGGGGACGCTCGACAACGCGCTCTACCCGGTGCTCTTCCTCGACTACCTCCGC
lat7-1  TGGGCGTCGGGGACGCTCGACAACGCGCTCTACCCGGTGCTCTTCCTCGACTACCTCCGC
lat7-2  TGGGCGTCGGGGACGCTCGACAACGCGCTCTACCCGGTGCTCTTCCTCGACTACCTCCGC

        ....:..370....:..380....:..390....:..400....:..410....:..420
LAT7    TCCGGCGGGGGGCTCGTGCTCTCCCCGCCGGCCCGCTCCCTCGCCGTGCTCGCGCTCACC
lat7-1  TCCGGCGGGGGGCTCGTGCTCTCCCCGCCGGCCCGCTCCCTCGCCGTGCTCGCGCTCACC
lat7-2  TCCGGCGGGGGGCTCGTGCTCTCCCCGCCGGCCCGCTCCCTCGCCGTGCTCGCGCTCACC

        ....:..430....:..440....:..450....:..460....:..470....:..480
LAT7    GCCGCGCTCACCTACCTCAACTTCCGGGGGCTCCACCTCGTCGGCCTCTCCGCGCTGGCG
lat7-1  GCCGCGCTCACCTACCTCAACTTCCGGGGGCTCCACCTCGTCGGCCTCTCCGCGCTGGCG
lat7-2  GCCGCGCTCACCTACCTCAACTTCCGGGGGCTCCACCTCGTCGGCCTCTCCGCGCTGGCG
```

```
        ....:..490....:..500....:..510....:..520....:..530....:..540
LAT7    CTCACCGCGTTCTCGCTCTCCCCGTTCGTCGCGCTCGCCGTGCTCGCCGCCCCCAAGATC
lat7-1  CTCACCGCGTTCTCGCTCTCCCCGTTCGTCGCGCTCGCCGTGCTCGCCGCCCCCAAGATC
lat7-2  CTCACCGCGTTCTCGCTCTCCCCGTTCGTCGCGCTCGCCGTGCTCGCCGCCCCCAAGATC

        ....:..550....:..560....:..570....:..580....:..590....:..600
LAT7    CGCCCGTCGCGGTGGCTCGCCGTGAACGTGGCCGCCGTTGAGCCGCGCGCCTACTTCAAC
lat7-1  CGCCCGTCGCGGTGGCTCGCCGTGAACGTGGCCGCCGTTGAGCCGCGCGCCTACTTCAAC
lat7-2  CGCCCGTCGCGGTGGCTCGCCGTGAACGTGGCCGCCGTTGAGCCGCGCGCCTACTTCAAC

        ....:..610....:..620....:..630....:..640....:..650....:..660
LAT7    TCCATGTTCTGGAACCTCAACTACTGGGACAAGGCGAGCACGCTTGCCGGCGAGGTGGAG
lat7-1  TCCATGTTCTGGAACCTCAACTACTGGGACAAGGCGAGCACGCTTGCCGGCGAGGTGGAG
lat7-2  TCCATGTTCTGGAACCTCAACTACTGGGACAAGGCGAGCACGCTTGCCGGCGAGGTGGAG

        ....:..670....:..680....:..690....:..700....:..710....:..720
LAT7    GAGCCGAGGAAGACGTTCCCGAAGGCGGTGTTCGGCGCGGTGGGGCTCGTCGTGGGCGCG
lat7-1  GAGCCGAGGAAGACGTTCCCGAAGGCGGTGTTCGGCGCGGTGGGGCTCGTCGTGGGCGCG
lat7-2  GAGCCGAGGAAGACGTTCCCGAAGGCGGTGTTCGGCGCGGTGGGGCTCGTCGTGGGCGCG

        ....:..730....:..740....:..750....:..760....:..770....:..780
LAT7    TACCTCATCCCGCTCCTCGCCGGGACGGGCGCGCTGCCGTCGGAGACGGCGGGGGAGTGG
lat7-1  TACCTCATCCCGCTCCTCGCCGGGACGGGCGCGCTGCCGTCGGAGACGGCGGGGGAGTGG
lat7-2  TACCTCATCCCGCTCCTCGCCGGGACGGGCGCGCTGCCGTCGGAGACGGCGGGGGAGTGG

        ....:..790....:..800....:..810....:..820....:..830....:..840
LAT7    ACGGACGGGTTCTTCTCCGTGGTCGGCGACCGGATCGGCGGGCCGTGGCTGCGCGTGTGG
lat7-1  ACGGACGGGTTCTTCTCCGTGGTCGGCGACCGGATCGGCGGGCCGTGGCTGCGCGTGTGG
lat7-2  ACGGACGGGTTCTTCTCCGTGGTCGGCGACCGGATCGGCGGGCCGTGGCTGCGCGTGTGG

        ....:..850....:..860....:..870....:..880....:..890....:..900
LAT7    ATCCAGGCCGCCGCGGCCATGTCCAACATGGGGCTCTTCGAGGCCGAGATGAGCGGCGAC
lat7-1  ATCCAGGCCGCCGCGGCCATGTCCAACATGGGGCTCTTCGAGGCCGAGATGAGCGGCGAC
lat7-2  ATCCAGGCCGCCGCGGCCATGTCCAACATGGGGCTCTTCGAGGCCGAGATGAGCGGCGAC

        ....:..910....:..920....:..930....:..940....:..950....:..960
LAT7    TCGTTCCAGCTCCTCGGCATGGCGGAGATGGGCATGATCCCGGCGATCTTCGCGCGCAGG
lat7-1  TCGTTCCAGCTCCTCGGCATGGCGGAGATGGGCATGATCCCGGCGATCTTCGCGCGCAGG
lat7-2  TCGTTCCAGCTCCTCGGCATGGCGGAGATGGGCATGATCCCGGCGATCTTCGCGCGCAGG

        ....:..970....:..980....:..990....:.1000....:.1010....:.1020
LAT7    TCGCGCCACGGCACGCCGACGTACAGCATCCTCTGCTCGGCCACCGGCGTCGTCATCCTC
lat7-1  TCGCGCCACGGCACGCCGACGTACAGCATCCTCTGCTCGGCCACCGGCGTCGTCATCCTC
lat7-2  TCGCGCCACGGCACGCCGACGTACAGCATCCTCTGCTCGGCCACCGGCGTCGTCATCCTC

        ....:.1030....:.1040....:.1050....:.1060....:.1070....:.1080
LAT7    TCCTTCATGAGCTTCCAGGAGATCGTCGAGTTCCTCAACTTCCTCTACGGCCTCGGGATG
lat7-1  TCCTTCATGAGCTTCCAGGAGATCGTCGAGTTCCTCAACTTCCTCTACGGCCTCGGGATG
lat7-2  TCCTTCATGAGCTTCCAGGAGATCGTCGAGTTCCTCAACTTCCTCTACGGCCTCGGGATG

        ....:.1090....:.1100....:.1110....:.1120....:.1130....:.1140
LAT7    CTCGCCGTGTTCGCCGCCTTCGTCAAGCTCCGCGTCAAGGACCCCGACCTCCCCCGCCCG
lat7-1  CTCGCCGTGTTCGCCGCCTTCGTCAAGCTCCGCGTCAAGGACCCCGACCTCCCCCGCCCG
lat7-2  CTCGCCGTGTTCGCCGCCTTCGTCAAGCTCCGCGTCAAGGACCCCGACCTCCCCCGCCCG
```

```
        ....:.1150....:.1160....:.1170....:.1180....:.1190....:.1200
LAT7    TACCGGATCCCCGTCGGCGCCGCGGGCGCCGCCGCCATGTGCGTCCCGCCCGTCGTCCTC
lat7-1  TACCGGATCCCCGTCGGCGCCGCGGGCGCCGCCGCCATGTGCGTCCCGCCCGTCGTCCTC
lat7-2  TACCGGATCCCCGTCGGCGCCGCGGGCGCCGCCGCCATGTGCGTCCCGCCCGTCGTCCTC

        ....:.1210....:.1220....:.1230....:.1240....:.1250....:.1260
LAT7    ATCACCACCGTCATGTGCCTCGCCTCCGCCAGGACGCTCGTCGTCAGCGCCGCCGTGGCC
lat7-1  ATCACCACCGTCATGTGCCTCGCCTCCGCCAGGACGCTCGTCGTCAGCGCCGCCGTGGCC
lat7-2  ATCACCACCGTCATGTGCCTCGCCTCCGCCAGGACGCTCGTCGTCAGCGCCGCCGTGGCC

        ....:.1270....:.1280....:.1290....:.1300....:.1310....:.1320
LAT7    GTCGCCGGCGTCGCCATGTACTACGGCGTCGAGCACATGAAGGCCACCGGCTGCGTCGAG
lat7-1  GTCGCCGGCGTCGCCATGTACTACGGCGTCGAGCACATGAAGGCCACCGGCTGCGTCGAG
lat7-2  GTCGCCGGCGTCGCCATGTACTACGGCGTCGAGCACATGAAGGCCACCGGCTGCGTCGAG

        ....:.1330....:.1340....:.1350....:.1360....:.1370....:.1380
LAT7    TTCTTGACGCCGGTGCCGCCTGACAGCCTCCGTGGATCATCATCATCATCATCCTCATCG
lat7-1  TTCTTGACGCCGGTGCCGCCTGACAGCCTCCGTGGATCATCATCATCATCATCCTCATCG
lat7-2  TTCTTGACGCCGGTGCCGCCTGACAGCCTCCGTGGATCATCATCATCATCATCCTCATCG

        ....:.1390....:.1400....:.1410....:.1420....:.1430....:.1440
LAT7    GCAGCGTCCGACAACGGCGGCGACGACGACGTCGAGGACGTCTGCGCCCTCCTCCTCGCC
lat7-1  GCAGCGTCCGACAACGGCGGCGACGACGACGTCGAGGACGTCTGCGCCCTCCTCCTCGCC
lat7-2  GCAGCGTCCGACAACGGCGGCGACGACGACGTCGAGGACGTCTGCGCCCTCCTCCTCGCC

        ....:.1450....:.1460....:.1470....:.1480....:.1490.
LAT7    GCCGGCGAGCACGCCGGAGAAGGCGTCAGTGTCAGCAAGGAGAATTATTAG
lat7-1  GCCGGCGAGCACGCCGGAGAAGGCGTCAGTGTCAGCAAGGAGAATTATTAG
lat7-2  GCCGGCGAGCACGCCGGAGAAGGCGTCAGTGTCAGCAAGGAGAATTATTAG
```

**S6 Appendix. Putative rice lat1, lat5, and lat7 protein sequences from single mutant lines.**

Putative lat1-1 protein sequence:
MADTGGRPEVSLATVRSPGHPAASTTAAAAADLGHADTGQEKPTVESAQPANGAAPMGECGTEYRGLPDGDAGGP
MPSSARTVSMIPLIFLIFYEVSGGPFGIEDSVGAAGPLLAIIGFLVLPVIWSIPEALITAELGAMFPENGGYVVW
VASALGPYWGFQQGWMKWLSGVIDNALYPVLFLDYLKSGVPALGGGAPRAFAVVGLTAVLTLLNYRGLTVVGWVA
ICLGVFSLLPFFVMGLIALPKLRPARWLVIDLHNVDWNLYLNTLFWNLNYWDSISTLAGEVKNPGKTLPKALFYA
VIFVVVAYLYPLLAGTGAVPLDRGQWTDGYFADIAKLLGGAWLMWWVQSAAALSNMGMFVAEMSSDSYAGHGGAG
HAPVLLRGAVAVRHAAGGHPLLGLRRAAALDDELPGDRGGRELPLLLRHAPRVRRLHPAPGEAPRRGAPIQGAAG
HSRVRGDAGAADGADRRGARAVHAEGGGGEPRRGGHGARAAAGAEVRGEEAVAEVLR*

Putative lat1-2 protein sequence:
MADTGGRPEVSLATVRSPGHPAASTTAAAAADLGHADTGQEKPTVESAQPANGAAPMGECGTEYRGLPDGDAGGP
MPSSARTVSMIPLIFLIFYEVSGGPFGIEDSVGAAGPLLAIIGFLVLPVIWSIPEALITAELGAMFPENGGYVVW
VASALGPYWGFQQGWMKWLSGVIDNALYPVLFLDYLKSGVPALGGGAPRAFAVVGLTAVLTLLNYRGLTVVGWVA
ICLGVFSLLPFFVMGLIALPKLRPARWLVIDLHNVDWNLYLNTLFWNLNYWDSISTLAGEVKNPGKTLPKALFYA
VIFVVVAYLYPLLAGTGAVPLDRGQWTDGYFADIAKLLGGAWLMWWVQSAAALSNMGMFVAEMSSDSYQLAGHGG
AGHAPVLLRGAVAVRHAAGGHPLLGLRRAAALDDELPGDRGGRELPLLLRHAPRVRRLHPAPGEAPRRGAPIQGA
AGHSRVRGDAGAADGADRRGARAVHAEGGGGEPRRGGHGARAAAGAEVRGEEAVAEVLR*

Putative lat5-1 protein sequence:
MTNAWISPSVVALCPSPLPSSRLPGSVLSCWPDSRGIRRGAGEGTAGQTLRPARGFTVEKLRNTAITRANSACLP
MEDCVGIKYSSVNEGEERKGAMASQRFPSSHSFSSYSMKFLGVRLGLRIVSRLLAHS*

Putative lat5-2 protein sequence:
MTNAWISPSVVALCPSPLPSSRLPGSVLSCWPDSRGIRRGAGEGTAGQTLRPARGFTVEKLRNTAITRANSACLP
MEDCVGIKYSSVNEGEERKGASQRFPSSHSFSSYSMKFLGVRLGLRIVSRLLAHS*

Putative lat7-1 protein sequence:
MTGACEAAPARRRGLTVLPLVALIFYDVSGGPFGIEDSVRAGGGALLPILGFLVLPVLWSLPRRSSPPSSPPRSP
PTPATSPGSPPRSAPPRRSSSGSPSGRRGRSTTRSTRCSSSTTSAPAGGSCSPRRPAPSPCSRSPPRSPTSTSGG
STSSASPRWRSPRSRSPRSSRSPCSPPPRSARRGGSP*

Putative lat7-2 protein sequence:
MTGACEAAPARRRGLTVLPLVALIFYDVSGGPFGIEDSVRAGGGALLPILGFLVLPVLWSPRRSSPPSSPPRSPP
TPATSPGSPPRSAPPRRSSSGSPSGRRGRSTTRSTRCSSSTTSAPAGGSCSPRRPAPSPCSRSPPRSPTSTSGGS
TSSASPRWRSPRSRSPRSSRSPCSPPPRSARRGGSP*
